# Myosin VI orchestrates estrogen-driven gene expression in breast cancer cells

**DOI:** 10.1101/2025.11.13.688007

**Authors:** Yukti Hari-Gupta, Danielle Lambert, Isabel W. Shahid-Fuente, Natalia Fili, Ália dos Santos, Alexandria Sprules, Hyejeong Rosemary Kim, Alexander W. Cook, Peter J.I. Ellis, Jesse Aaron, Teng-Leong Chew, Lin Wang, Christopher P. Toseland

## Abstract

Approximately 75% of breast cancer cases are estrogen receptor (ER)-positive, with endocrine therapies forming the foundation of treatment. However, therapeutic resistance remains a major clinical challenge, necessitating the identification of new molecular targets. Myosin VI (MVI), a motor protein increasingly linked to cancer, directly interacts with the ER and plays a key role in the spatial regulation of RNA Polymerase II. Notably, expression levels of MVI and ER are positively correlated across breast cancer tissues. Here, we present a multidisciplinary investigation combining advanced imaging, genomic profiling, and phenotypic characterisation to elucidate the interplay between MVI and the ER. We demonstrate that MVI nuclear localisation is dynamically regulated by estrogen signalling and ER activity. Conversely, ER nuclear localisation requires active MVI, suggesting a reciprocal regulatory mechanism. Furthermore, MVI influences subnuclear architecture, modulating ER transcriptional activity and downstream gene expression programmes that govern cell proliferation and migration. Importantly, pharmacological inhibition of MVI enhances the effect of hormone therapy, resulting in greater disruption of ER function than monotherapy alone. Moreover, MVI inhibition also suppresses the activity of hormone-resistant ER mutants, highlighting its potential to overcome therapy resistance. Our findings establish MVI as a critical regulator of ER nuclear dynamics and gene expression, supporting its candidacy as a novel therapeutic target in ER-positive breast cancer.

## INTRODUCTION

Breast cancer remains a leading cause of mortality and morbidity worldwide ^1^, and approximately 75% of all breast cancers are classified as estrogen receptor (ER)-positive ^2^. ER-positive tumours require, and respond to, hormones such as estrogen to drive gene expression and disease progression through ligand-dependent actions of the ER. In the canonical mechanism, estrogen binds the ER to form an active complex that translocates to the nucleus. There, the ER recruits co-regulator proteins ^3^ and binds to estrogen response elements (EREs) in target gene promoters, driving transcription. Hormone therapy, which has been shown to increase recurrence-free survival and overall survival, is given to patients with ER-positive tumours. Several classes of hormone therapy have been developed, each aimed at disrupting different pathway components of ER signalling, to ultimately block the interaction between receptor and hormone ^4^. Firstly, Selective Estrogen Receptor Modulators (SERMs) compete with estrogen to bind the ER, to promote a conformational change so that it cannot bind to its target genes ^5,6^. The second class are Selective ER Degraders (SERDs). These also compete with estrogen to bind to the ER, although this is with greater affinity compared to SERMs. Moreover, SERDs promote proteasomal degradation of the ER, reducing the levels available for estrogen activation ^7^. Finally, Aromatase Inhibitors (AIs) target estrogen production thus preventing ligand-dependent ER activation and subsequent gene expression ^8^.

Unfortunately, approximately 50% of patients on endocrine therapy develop resistance ^9–11^, leading to tumour progression and, in the absence of effective second-line treatments, ultimately death. This frequency of resistance underlines the need for novel combination or second-line therapies to improve patient outcomes. Therefore, it is crucial research focuses on the development of secondary, or combinatory, treatments to improve patient outcomes.

Myosin VI (MVI) is an actin-based motor protein which has been noted to be highly expressed in many cancers including breast and prostate (Figure 1), where it has been shown to play roles in regulating cell proliferation and migration ^12,13^. MVI has diverse cytoplasmic and nuclear functions ranging from endocytosis through to transcription ^14^. Within the nucleus, MVI has been shown to have roles in transcription via binding of RNA polymerase II complexes and aiding its spatial organisation ^15–17^. In this manner, MVI physically stabilises RNAPII at sites of transcription. Notably, MVI can directly bind nuclear hormone receptors (including ER) via a C-terminal LxxLL motif, analogous to classic co-activators ^15^. We propose that this MVI–ER interaction helps recruit MVI to specific ER-regulated genes, thereby positioning MVI to regulate transcription at those sites.

**Figure 1.**
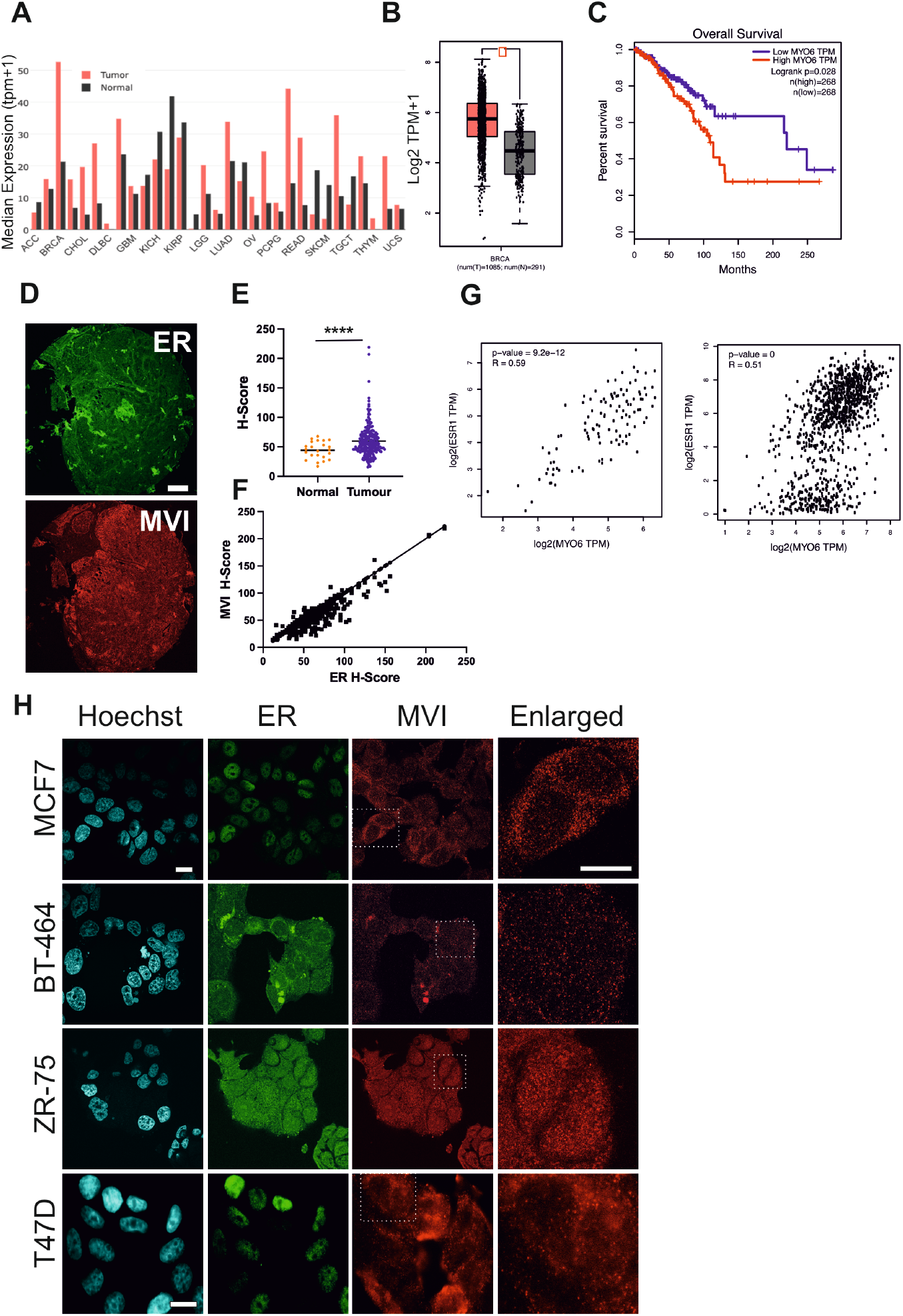
Expression of myosin VI in breast cancer and its association with the Estrogen receptor. (A) *MYO6* expression across different cancer types. The bar height represents the median expression (Log_2_ Transcripts per million+1) of certain tumour type (red) or normal tissue (black). (B) Expression level (Transcripts per million) of *MYO6* in breast cancer vs normal breast tissue. (C) Kaplan-Meier plots of overall breast cancer survival with low (n =2 68) and high (n = 268) MYO6 expression. (D) Representative example of tissue array with Immunofluorescence staining against MVI (red) and ER (green). Scale bar = 150 µm. (E) MVI H-Score calculated from three independent tissue arrays. Normal (n = 21) Tumour (n = 209). ****p <0.0001 by unpaired t-test. (F) Correlation analysis for MVI and ER H-Scores from three independent tumour tissue arrays (n=209). Correlation coefficient 0.78. (G) *MYO6* and *ESR1* transcript expression correlation in normal breast (TCGA and GTex databases) and breast cancer tissue (TCGA database), respectively. (H) Representative widefield Immunofluorescence staining against MVI (red), ER (green) and DNA (cyan) in the stated cells (Scale bar 10 μm).

In this work, we investigate MVI as a critical co-regulator of ER-driven gene expression. Our findings reveal that disrupting MVI not only impairs normal estrogen-dependent transcription and tumour cell proliferation but also enhances the efficacy of endocrine therapies and overcomes resistance caused by constitutively active ER mutants. These insights establish MVI as a promising new therapeutic target in ER-positive breast cancer.

## RESULTS

### The expression levels of myosin VI and estrogen receptor are correlated across breast cancer tissues

Evaluation of The Cancer Genome Atlas (TCGA) and Genotype Tissue Expression (GTEx) databases through the Gene Expression Profiling Interactive Analysis (GEPIA) ^18^ server revealed that MVI is significantly up regulated in many cancers (19/29) (Figure 1A), including STAD, READ, PRAD and BRCA (P<0.05) (Figure 1B). Conversely, some cancers have a down-regulation of MVI (10/29). Exploring the expression level in breast cancer in more detail, MVI was significantly up-regulated in HER2, Luminal A and Luminal B types but not within basal type (Supplementary Figure 1A). Moreover, MVI expression level is not related to cancer stage (Supplementary Figure 1B). Lastly, high level expression of MVI in breast cancer impacts overall breast cancer survival rates (Figure 1C).

We have previously revealed that there is a direct interaction between MVI and the ER ^15^. To understand the nature of this interaction with respect to tumours, we assessed the expression of MVI and ER in breast tissue within a tissue array stained for MVI and ER (Figure 1D). Whole tissue samples were imaged and H-scores calculated to determine protein levels. Staining intensity for MVI was enhanced within the breast cancer samples (Figure 1E). Correlation analysis showed a strong positive correlation (Figure 1F, r=0.7835, P=<0.0001), where high levels of ER were associated with high MVI levels in cancer tissue. The correlated protein expression level was also consistent with transcript expression level in normal and tumour samples from GTEx and TCGA databases (Figure 1G).

Exploring the MVI expression in more detail, there is a significantly higher usage of the non-insert isoform of MVI (Supplementary Figure 1C and 1D), noted as MYO6-003. This isoform is the one we previously identified as acting within the cell nucleus to support gene expression ^15^. Consistent with our previous findings, MVI is expressed and found within the nucleus of breast cancer cell lines (Figure 1H) but the level does vary across the cell lines.

### The myosin VI and estrogen receptor interaction is required for nuclear localisation

The clinical data raise questions regarding the interplay between MVI and the ER which are known to interact. To this end, we examined whether manipulating hormones or protein levels impacts the nuclear localisation of each protein.

We measured the nuclear population of MVI, whilst directly and indirectly manipulating the ER. Firstly, following 48 hrs starvation of the cells of hormone (Figure 2A), we observed an expected decrease in the ER nuclear levels (Figure 2B and 2C), as well as a decrease in the MVI level (Figure 2D). Conversely, stimulation with estradiol led to an increase in ER and MVI within the nucleus (Figure 2B-D). In addition, knockdown of ER led to a significant decrease in nuclear MVI.

**Figure 2.**
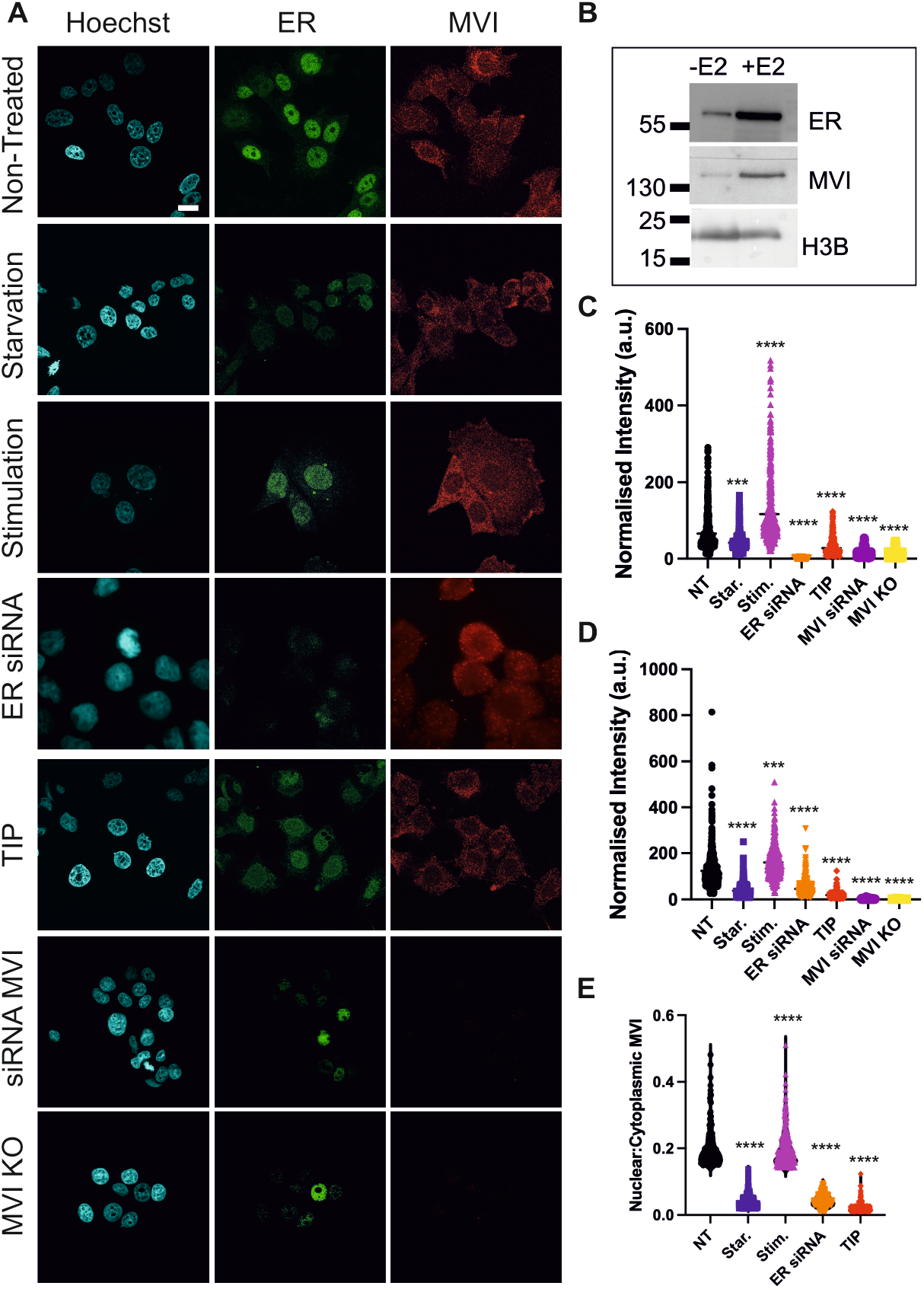
The interdependency of Myosin VI and ER nuclear localisation. (A) Representative Immunofluorescence staining against MVI (red), ER (green) and DNA (cyan) in MCF7 cells (Scale bar 10 μm). Hormone starvation occurred for 48 hrs and stimulation was performed upon starved cells for 24 hr with 10 nM estradiol. siRNA knockdowns were performed for 48 hrs. 25 µM TIP for 6 hrs. Western blots for knockdown and knockout (KO) conditions are shown in Supplementary Figure 2. (B) Western-blot of the nuclear fraction against MVI and ER under hormone starvation and stimulation conditions in MCF7 cells. (C) and (D) Quantification of nuclear ER and MVI normalised intensity across the experimental conditions. n=560 cells per condition across three independent experiments. Quantification of control DMSO and scrambled siRNA is shown in Supplementary Figure 2. ***p <0.001 ****p <0.0001 by unpaired t-test compared to non-treated (NT) conditions. (E) Quantification of the nuclear:cytoplasmic ratio for MVI data measured in (D). ****p <0.0001 by unpaired t-test compared to non-treated conditions, except for stimulation which is compared to starvation conditions.

For all measurements, not only did the MVI intensity vary across the conditions, but the overall cellular distribution of MVI altered across the conditions, as identified in the nuclear:cytoplasmic ratio measurements (Figure 2E).

Having revealed the impact of ER activity on the localisation of nuclear MVI, we then explored the impact of MVI on ER localisation. Interestingly, siRNA transient knockdown of MVI, CRISPR-Cas9 MVI knockout, or motor inhibition through treatment with TIP, led to a significant decrease in the nuclear level of ER (Figure 2C). TIP also leads to the decrease in MVI nuclear levels (Figure 2D), consistent with our previous findings ^16^.

### The sub-nuclear organisation of the estrogen receptor is regulated by active myosin VI

Having established an interdependence in their nuclear localization, we next asked whether MVI influences the spatial organization of ER within the nucleus. To this end, we used Stochastic Optical Reconstruction Microscopy (STORM) and cluster analysis to measure the spatial organisation of MVI and ER in the nucleus with high precision (Figure 3A). Protein clustering is often related to the molecular function of a protein and is particularly important in the enhancement of enzymatic processes such as transcription. Clusters are defined as non-random distribution of proteins in relation to each other (Figure 3B). Specifically, we defined clusters as regions where a minimum of 5 neighbouring molecules are spaced at a distance smaller than the mean value of localisation precision from STORM acquisition (described in Methods) ^19^.

**Figure 3.**
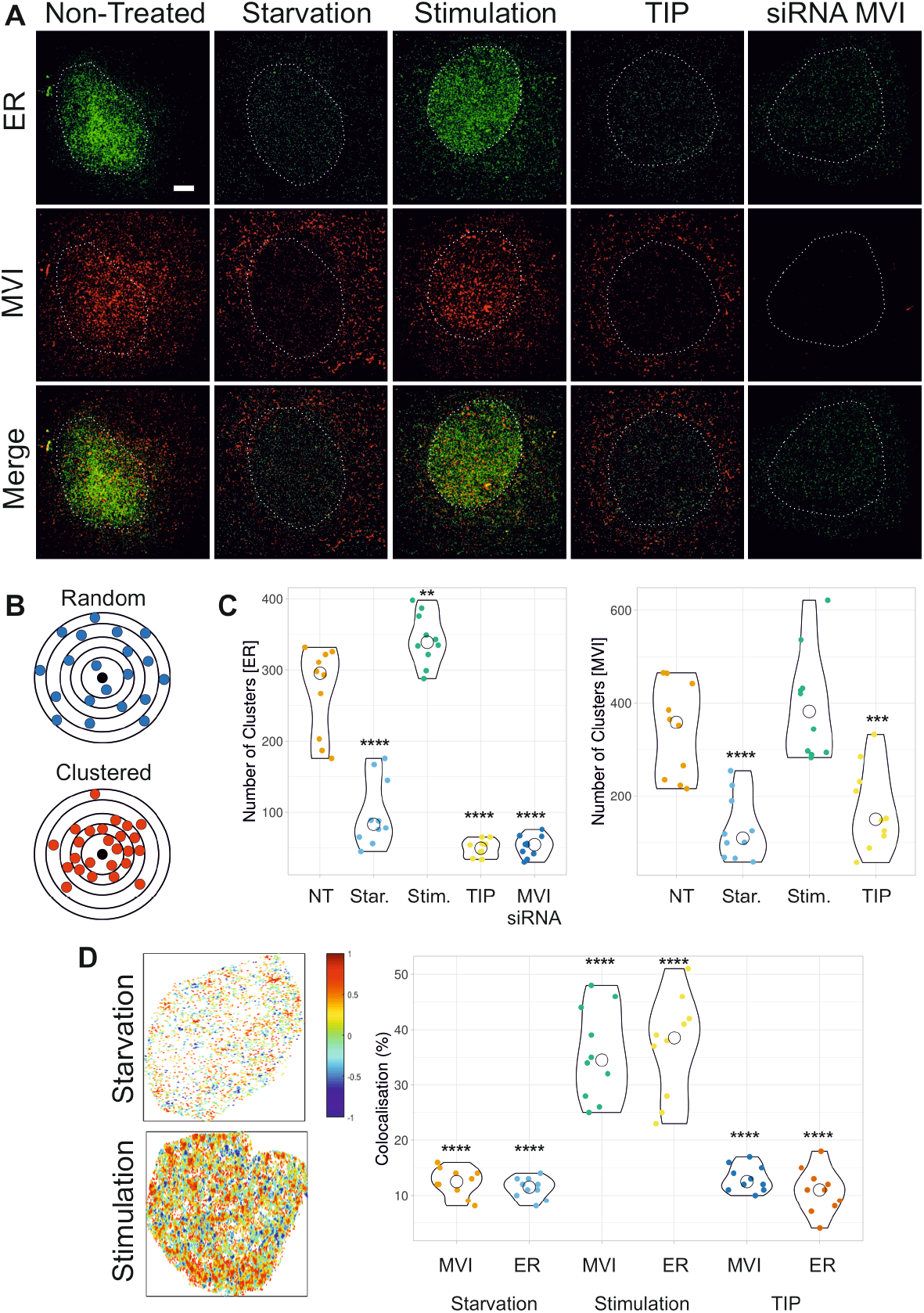
Nuclear organisation of Myosin VI and ER. A) Example STORM render images of MVI and ER under non-treated (NT), hormone starvation, estradiol stimulation, TIP and MVI knockdown (siRNA MVI) conditions (scale bar 2 □m). Hormone starvation occurred for 48 hrs and stimulation was performed upon starved cells for 24 hr with 10 nM estradiol. siRNA knockdowns were performed for 48 hrs. 25 µM TIP for 6 hrs. The white dotted lines represent a region of interest (ROI) containing the whole nucleus which was taken forward for cluster analysis. (B) Depiction of molecular clustering and random distribution of proteins where the core point (black dots) have ≥ 4 neighbours within a 1-unit radius (black circles) in the clustered distribution. (C) Cluster analysis of MVI and ER nuclear organisation. Clusters are defined by detecting a minimum of 5 molecules within a search area corresponding to the STORM localisation precision. The search area then propagates and a group of molecules is considered to be a cluster if at least 10 molecules are found. Individual data points correspond to the average value for a cell ROI (n = 10). The values depicted by black circles represent the number of clusters from the ROIs for each condition. DMSO and control siRNA measurements are shown in Supplementary Figure 3. Only statistically significant changes are highlighted **p <0.01, ***p <0.001, ****p <0.0001 by unpaired t-test compared to normal conditions. (D) Colocalisation analysis of MVI and ER clusters under hormone starvation, stimulation and TIP-treated conditions. Inset is a representative cluster colocalisation heatmap whereby DoC values of 1 are perfectly colocalised and −1 are separated from each other. Individual data points represent the percentage of each protein which is colocalised based on DoC threshold of above 0.4 (n=10) (see Supplementary Figure 4 for example histograms). The depicted by black circles values represent the mean from the ROIs for each protein. ****p <0.0001 by unpaired t-test compared to normal conditions for each protein.

MVI formed clusters in the nucleus of MCF7 cells (Figure 3A), as previously characterised in HeLa cells ^16^. Interestingly, the MVI clustering parameters were dependent upon hormone, whereby starvation leads to a significant decrease in cluster number from 290 to 85 clusters per nucleus (Figure 3C) and cluster area from 3400 nm^2^ to 1400 nm^2^ (Supplementary Figure 3). Then, stimulation with estradiol caused a significant increase in the number of clusters to 330 above starvation, While STORM provided a static snapshot of MVI and ER clusters, we next examined the dynamic behaviour of these molecules in live cells. If MVI is anchoring ER at transcription sites, we would expect to see changes in diffusion behaviour when MVI or ER is perturbed. We performed aberration-corrected multi-focal microscope (acMFM) system ^22^ to simultaneously track single-molecules across nine focal planes in live-cells, covering 4 μm in the *z* axis and 20 × 20 μm in *xy*. In this way, we were able to observe and track the 3D dynamics of stably expressed Halo-tagged MVI and SNAP-tagged ER across the whole nucleus in live MCF7 cells (Figure 4A and Supplementary Figure 5). The 3D trajectories represent freely diffusing molecules, whereas the spatially confined trajectories likely represent molecules bound in complexes (Figure 4B), which we previously identified as clusters (Figure 3). We determined the diffusion constant for each track by measuring the Mean Squared Displacement (MSD) and then plotted the average diffusion and rescue in the size of MVI clusters to 3400 nm^2^.

**Figure 4.**
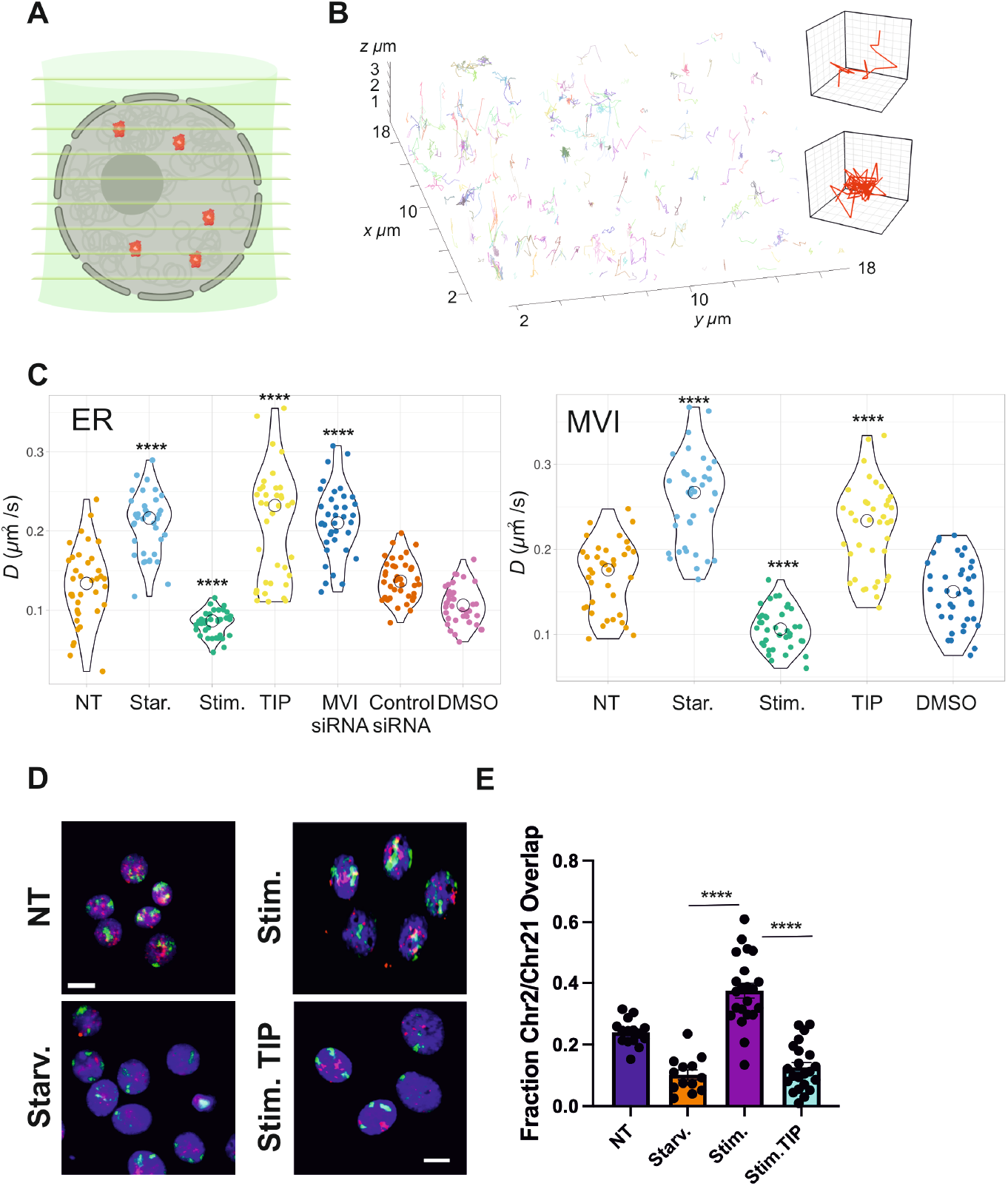
Live cell single molecule dynamics of Myosin VI and ER. (A) Cartoon depicting simultaneous acquisition of 9 focal planes covering 4 μm in the z-axisto perform live cell 3D single molecule tracking of fluorescently tagged proteins. (B) Example render of 3D single molecule trajectories. INSET: Representative trajectories of a diffusive (top inset) and a spatially confined molecule (bottom inset). (C) Plot of SNAP-ER and Halo-MVI diffusion constants under the stated conditions derived from fitting trajectories to an anomalous diffusion model, as described in methods. Individual data points correspond to the average value for a cell ROI (n = 38 from three independent experiments). Hormone starvation occurred for 48 hrs and stimulation was performed upon starved cells for 24 hr with 10 nM estradiol. siRNA knockdowns were performed for 48 hrs. 25 µM TIP for 6 hrs. The values represent the mean from the ROIs for each condition. Only statistically significant changes are highlighted ****p <0.0001 by unpaired t-test compared to normal conditions. (D) Nuclear myosin VI supports long-range chromatin reorganization. Chromosome PAINT of chromosome 2 (green) and 21 (magenta) in MCF7 cells under the stated conditions. Scale bar is 10 µm. (E) Fraction of chromosome 2 overlapping with chromosome 21 in non-treated (NT) MCF7 cells and in cells starved from hormone, stimulated with estradiol, and stimulation in the presence of TIP. Error bars represent SEM from three independent experiments. ****p <0.0001 by unpaired t-test compared to normal conditions.

**Figure 5.**
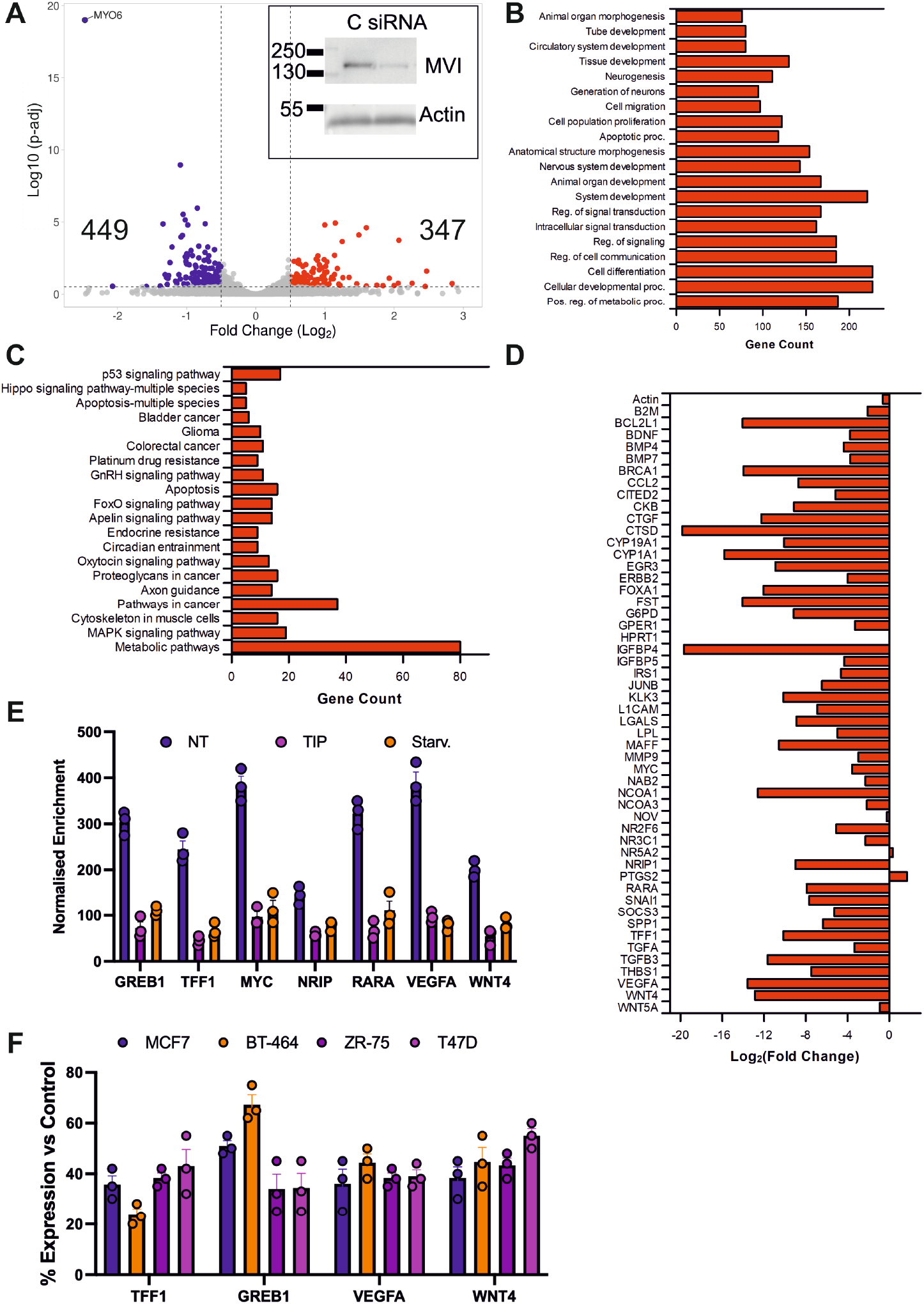
Myosin VI dependent changes in ER-driven gene expression. (A) Volcano plot of differentially expressed genes from RNA-seq following MVI siRNA knockdown in MCF7 cells. INSET: Western blot for the control “C” and MVI siRNA knockdown. (B) GO terms with gene count, for Biological Process corresponding to the differentially expressed genes following MVI knockdown. GO terms are plotted based on significant enrichment, as shown in Supplementary Table 1. (C) GO terms with gene count, for Biological Pathways corresponding to the differentially expressed genes following MVI knockdown. GO terms are plotted based on significant enrichment, as shown in Supplementary Table 1. (D) RT-qPCR Gene expression analysis of 28 ER regulated genes and housekeeping gene actin. MCF7 cells were hormone starved for 48 hrs then stimulated with 10 nM estradiol for 24 hrs in the presence and absence of 25 µM TIP. The plot depicts the Log_2_ fold change in expression between non-treated (NT) and TIP-treated conditions. (E) MVI ChIP-qPCR against labelled loci. Values are the average of three independent experiments. Cell lines were treated with 25 µM TIP for 6 hrs or starved from hormone for 48 hrs. Error bars represent SEM from three independent experiments. All conditions yielded statistically significant changes (p<0.0001) to non-treated (NT) condition by unpaired t-test. (F) RT-qPCR gene expression analysis of the stated genes within the four cell lines. Cell lines were treated with 25 µM TIP for 6 hrs. The plot depicts the percentage change in expression between non-treated and TIP-treated conditions. Error bars represent SEM from three independent experiments. TIP and DMSO controls for RT-qPCR are shown in Supplementary Figure 6. All conditions yielded statistically significant changes (p<0.0001) to control by unpaired t-test.

With respect to the ER, as expected there is a significant decrease in ER clusters and cluster area following hormone starvation, which was enhanced following hormone stimulation (Figure 3C and Supplementary Figure 3). Interestingly, TIP-treatment or siRNA knockdown of MVI also led to a decrease in ER clustering parameters, which was at a greater level than hormone starvation.

To evaluate a direct relationship between the proteins, the colocalisation between MVI and ER clusters was assessed. In order to quantify this colocalisation, we employed the Degree of Colocalisation (DoC) analysis ^20^ which employs a coordinate-based method to determine correlation between individual molecules in each channel ^21^. The colour-coded colocalisation cluster map reflects the DoC value for the ER and MVI. Applying a threshold of 0.4 allowed us to determine that 12% of MVI and 11% of ER molecules colocalise (Figure 3D and Supplementary Figure 4) under starvation conditions. This significantly rises upon hormone stimulation to 34% and 39% for MVI and ER, respectively. TIP treatment reduces the colocalisation level back to those observed under starvation. Overall, this suggests there is synergy between the populations of the two proteins. Nevertheless, there are significant pools of protein acting independently.

constant per cell (Figure 4C). Hormone starvation, or TIP treatment, led to an increase in mean MVI diffusion. Conversely, hormone stimulation led to a significant decrease in mean diffusion. These changes are consistent with hormone triggering localisation and clustering of MVI at sites of gene expression.

Consistent with the STORM measurements, hormone starvation, siRNA knockdown of MVI and TIP-treatment led to an increase in mean diffusion of SNAP-ER, whilst hormone stimulation led to a decrease in mean diffusion (Figure 4C). These results revealed that perturbation of MVI directly impacts the nuclear pool of ER, rather than solely shuttling protein from the cytoplasm to the nucleus.

Together, the STORM clustering data and the single-molecule tracking data consistently indicate that hormonal stimulation consolidates MVI and ER into slower-diffusing complexes, assumed to be transcription complexes, whereas perturbation of MVI leads to more diffuse ER. This would align with disruption of the ER’s engagement at transcription sites.

### The interdependency of myosin VI – ER functionality impacts large-scale chromatin architecture

The STORM and single molecule tracking data suggest that active MVI helps maintain the ER in a clustered state at gene expression sites. To test whether MVI activity affects large-scale chromatin architecture of ER-regulated genes, we performed FISH-based chromosome painting for two representative ER target genes (GREB1 on chromosome 2 and TFF1 on chromosome 21).

After transcription activation by estradiol, the spatial overlap between both chromosomes increased compared to a transcription-inhibited state under hormone starvation conditions (Figure 4D and 4E). Treatment with 25 μM TIP inhibited chromosome reorganization leading to spatial segregation of the chromosomes (Figure 4D and 4E). These chromosome PAINT measurements report territory-scale overlap and radial shifts consistent with hormonedependent nuclear remodelling. This assay does not resolve the 3D positions of *GREB1* or *TFF1*; therefore, conclusions about gene-level repositioning remain indirect.

### Myosin VI regulates ER-driven gene expression

To explore the global changes in gene expression which arise from the perturbations described above, we performed RNA-seq measurements under normal and MVI knockdown conditions. In total, we observed a significant change in the expression of 796 genes following knock-down. From this set, 347 genes were up-regulated and 449 were down-regulated (Figure 5A). With respect to the down regulated genes, promoter screening by the Eurkaryotic Promoter Database revealed that >80 percent have an ERE motif within their promoter.

The differentially expressed genes were taken forward to Gene Ontology (GO) analysis. The breakdown for GO Biological Process reveals that the majority of the processes affected are coupled to developmental, proliferation, migration and signalling (Figure 5B and Supplementary Table 1). Further exploration of the pathways (Figure 5C and Supplementary Table 2) reveals a significant number of cancer-associated pathways.

To validate and extend the RNA-seq findings, we next focused on known estrogen-responsive genes. We tested whether pharmacological MVI inhibition (TIP) alters their induction by estradiol, and whether MVI is physically present at their promoters. We performed hormone stimulation on MCF7 cells in the absence and presence of MVI inhibition by TIP (Figure 5D). We then used RT-qPCR to monitor the expression of estrogen-responsive genes, and actin as a control. In all but four genes, inhibition of MVI led to a decrease in gene expression.

Next, we probed a subset of those genes for MVI binding at promoters by Chromatin Immunoprecipitation (ChIP) (Figure 5E). MVI enrichment at promoter regions was significantly decreased in both TIP and hormone-starvation conditions. Finally, we performed RT-qPCR analysis on the same genes across MCF7, BT-464, ZR-75 and T47D cells in the presence and absence of TIP.

Inhibition is consistent across the genes and cell lines (Figure 5F).

The transcriptional profiling reinforces the imaging data whereby the physical clustering and colocalisation of MVI with ER correlates with functional gene activation. When MVI is perturbed, nuclear ER clusters disassemble and accordingly, the expression of ER-driven genes decreases.

### Myosin VI perturbation impedes cell growth and migration

The widespread transcriptional changes suggest that MVI perturbation might have significant cellular consequences. We therefore examined two phenotypes, proliferation and migration.

We performed live-cell growth assays under normal and MVI transient knockdown conditions in MCF7 and T47D cells. A three-fold decrease in growth was observed following transient knockdown (Figure 6A). Conversely, stable over-expression of MVI increased MCF7 cell growth four-fold under the same conditions (Figure 6B).

**Figure 6.**
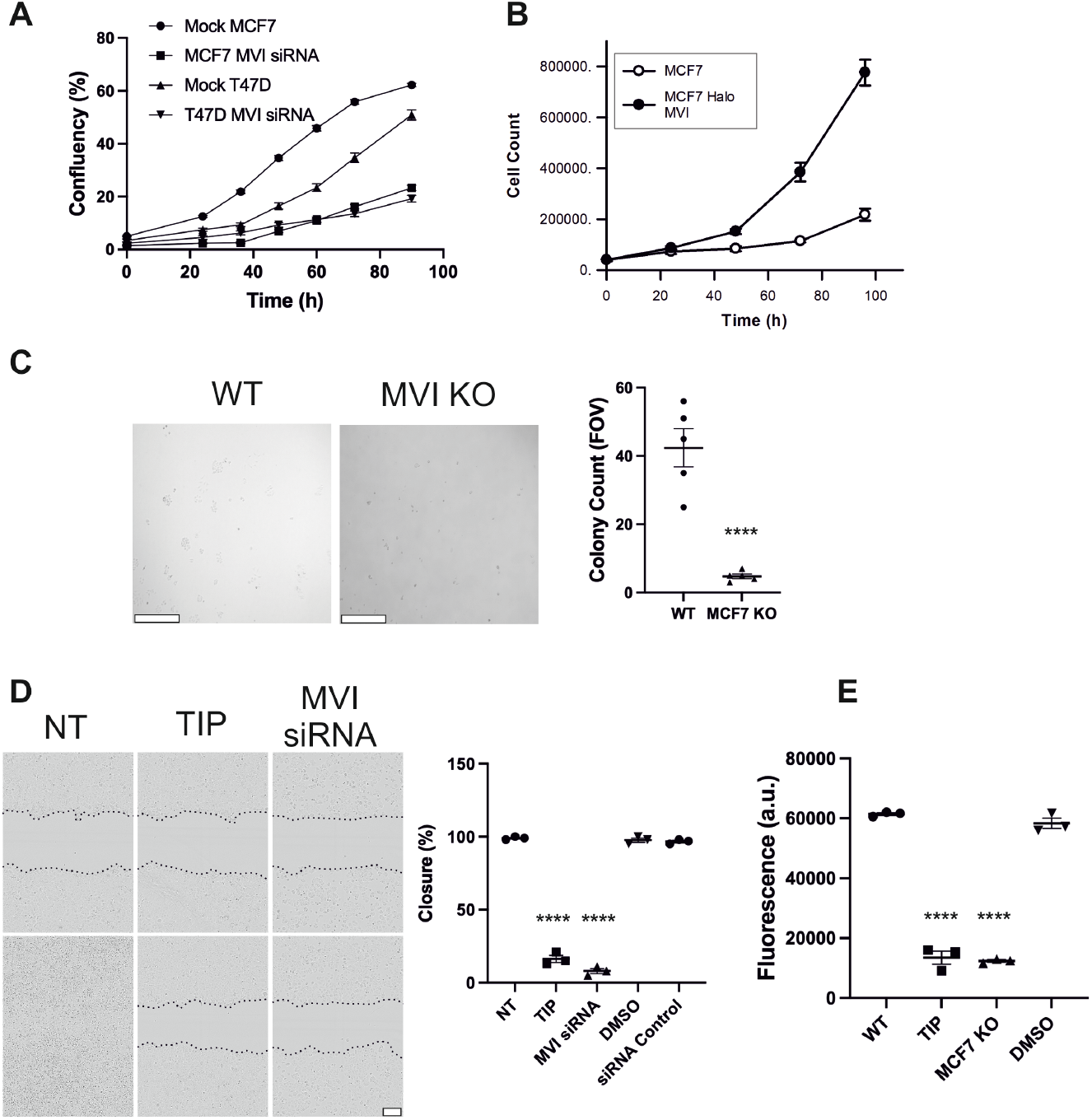
Myosin VI perturbation impacts cell growth and migration. (A) Real-time growth of MCF7 and T47D cells and corresponding measurements following MVI (MVI) siRNA knockdown for 24 hrs. Data represent three independent measurements and error bars show SEM. Western blots are shown in Supplementary Figure 7 for the MVI knockdown. (B) Time course of cell count for MCF7 and MCF7 Halo-MVI stable cell line. Data represent three independent measurements and error bars show SEM. (C) Colony formation assay for wild type MCF7 cells and MVI knockout MCF7 cells. Example images of the colonies are shown (Scale bar 600 µm). Data represent five independent measurements and error bars show SEM. Only statistically significant changes are highlighted ****p <0.0001 by unpaired t-test compared to normal conditions. (D) Scratch-wound healing assay for MCF7 cells under non-treated, treated with 25 µM TIP for 6 hrs and MVI knockdown for 24 hrs. Example images are shown at the start (top) and end of the measurement (bottom) (48 hrs). Data represent three independent measurements and error bars show SEM. Western blots are shown in Supplementary Figure 7 for the MVI knockdown. Only statistically significant changes are highlighted ****p <0.0001 by unpaired t-test compared to normal conditions. Scale bar 150 µm. (E) Tran-swell migration assay for wild type MCF7 cells, MVI knockout MCF7 cells and MCF7 cells treated with 25 µM TIP for 6 hrs. Data represent three independent measurements and error bars show SEM. Only statistically significant changes are highlighted ****p <0.0001 by unpaired t-test compared to normal conditions.

The growth defect was matched by the inability of MVI knockoutMCF7 cells to form colonies (Figure 6C). Overall, this change is indicative of a larger cellular response to the perturbation of the ER through MVI, and the resulting alterations in gene expression.

Along with growth related phenotypes, we also explored for migratory defects within MCF7 cells. Scratch wound healing assays were performed which revealed that TIP and siRNA knockdown of MVI significantly impeded wound closure (Figure 6D). Similarly, transwell migration assays were performed with TIP treated and MCF7 MVI knockout cells which revealed a significantly reduced migratory behaviour when MVI is perturbed (Figure 6E). We have also confirmed that inhibition of MVI is equivalent to transient knockdown, in terms of gene expression analysis (Supplementary Figure 7B). Moreover, the lack of enhanced impact of TIP upon siRNA treated cells identifies the MVI specific targeting of TIP.

**Figure 7.**
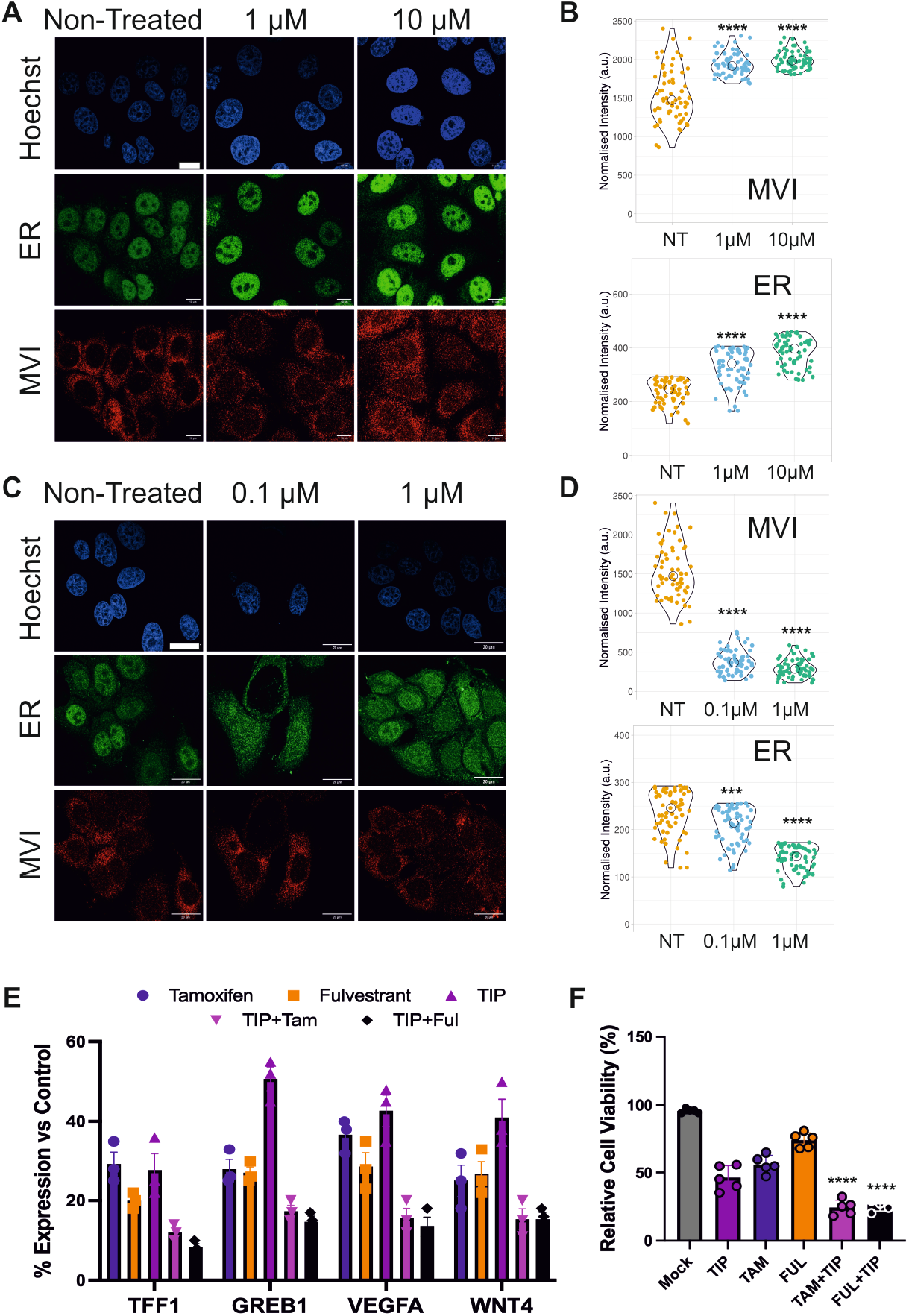
Inhibition of Myosin VI enhances endocrine therapy. (A) Representative Immunofluorescence staining against MVI (red), ER (green) and DNA (cyan) in MCF7 cells treated with the stated concentrations of tamoxifen for 6 hrs (Scale bar 10 μm). (B) Quantification of nuclear MVI and ER normalised intensity across the experimental conditions. n=66 cells per condition across three independent experiments. Quantification of control DMSO is shown in Supplementary Figure 2. ****p <0.0001 by unpaired t-test compared to normal conditions. (C) Representative Immunofluorescence staining against MVI (red), ER (green) and DNA (cyan) in MCF7 cells treated with the stated concentrations of fulvestrant for 6 hrs (Scale bar 20 μm). (D) Quantification of nuclear MVI and ER normalised intensity across the experimental conditions. n=66 cells per condition across three independent experiments. Quantification of control DMSO is shown in Supplementary Figure 2. ***p <0.001, ****p <0.0001 by unpaired t-test compared to normal conditions. (E) RT-qPCR gene expression analysis of the stated genes. Cells were treated with 1 µM Tamoxifen or 0.1 µM Fulvestrant in the absence and presence of 25 µM TIP for 6 hrs. The plot depicts the percentage change in expression between non-treated and treated cells. Error bars represent SEM from three independent experiments. DMSO controls for RT-qPCR are shown in Supplementary Figure 6. All conditions yielded statistically significant changes (p<0.0001) to control by unpaired t-test. (F) MTT cell viability assay under the same conditions as RT-qPCR. All data are normalised to non-treated control. ****p <0.0001 by unpaired t-test compared to single treatment condition.

### Targeting myosin VI in combination with endocrine therapy is more effective than monotherapy

Given the functional interplay between MVI and ER, we next explored how standard endocrine therapies influence MVI, and whether dual targeting of ER and MVI could be more effective. We examined the effects of a SERM (tamoxifen Figure 7A and 7B) and a SERD (fulvestrant Figure 7C and 7D) on MVI’s nuclear localization. As expected, these drugs alter ER levels/localization in a concentration-dependent manner. Importantly, we found they had corresponding effects on MVI localization, reinforcing the functional link between the two proteins.

This reciprocal behaviour prompted us to test a combination treatment hypothesis that inhibiting MVI could enhance the efficacy of ER-directed therapy. To this end, the lowest concentration of each therapy was combined with 25 µM TIP over 6 hours. The effect of the monotherapies and the combination therapy was assessed on the expression of representative ER-responsive genes. In all cases, the combination led to a significant decrease in gene expression compared to individual treatments (Figure 7E). Lastly, we monitored cell viability and once again found that combined treatment caused a significant decrease in viability over single treatment (Figure 7F). We therefore conclude that targeting the ER and MVI leads to a greater efficiency for inhibiting the activity of the ER.

### Myosin VI inhibition overcomes endocrine resistant ER mutants

A major clinical challenge arises from mutations in the ER which renders tumours estrogen-independent and resistant to therapies like tamoxifen or fulvestrant. We therefore asked whether MVI is still required for the activity of such mutant ER proteins.

To model hormone-independent ER activity, we expressed two common constitutively active ERα mutants (Y537S and D538G) in MCF7 cells. Remarkably, even though these mutants drive high target gene expression in the absence of hormone, the inhibition of MVI (with TIP) reduced their target gene expression to near baseline (Figure 8A– B). Moreover, TIP treatment of transfected cells decreased their viability (Figure 8C). Quantification of ER nuclear levels was not performed due to the high protein levels arising from the transient over expression. Qualitatively, nuclear levels of the mutant ER proteins were still present which suggests MVI disruption occurs within the nucleus. Taken together, MVI represents a potential therapeutic target to overcome endocrine therapy resistance caused by ER mutations.

**Figure 8.**
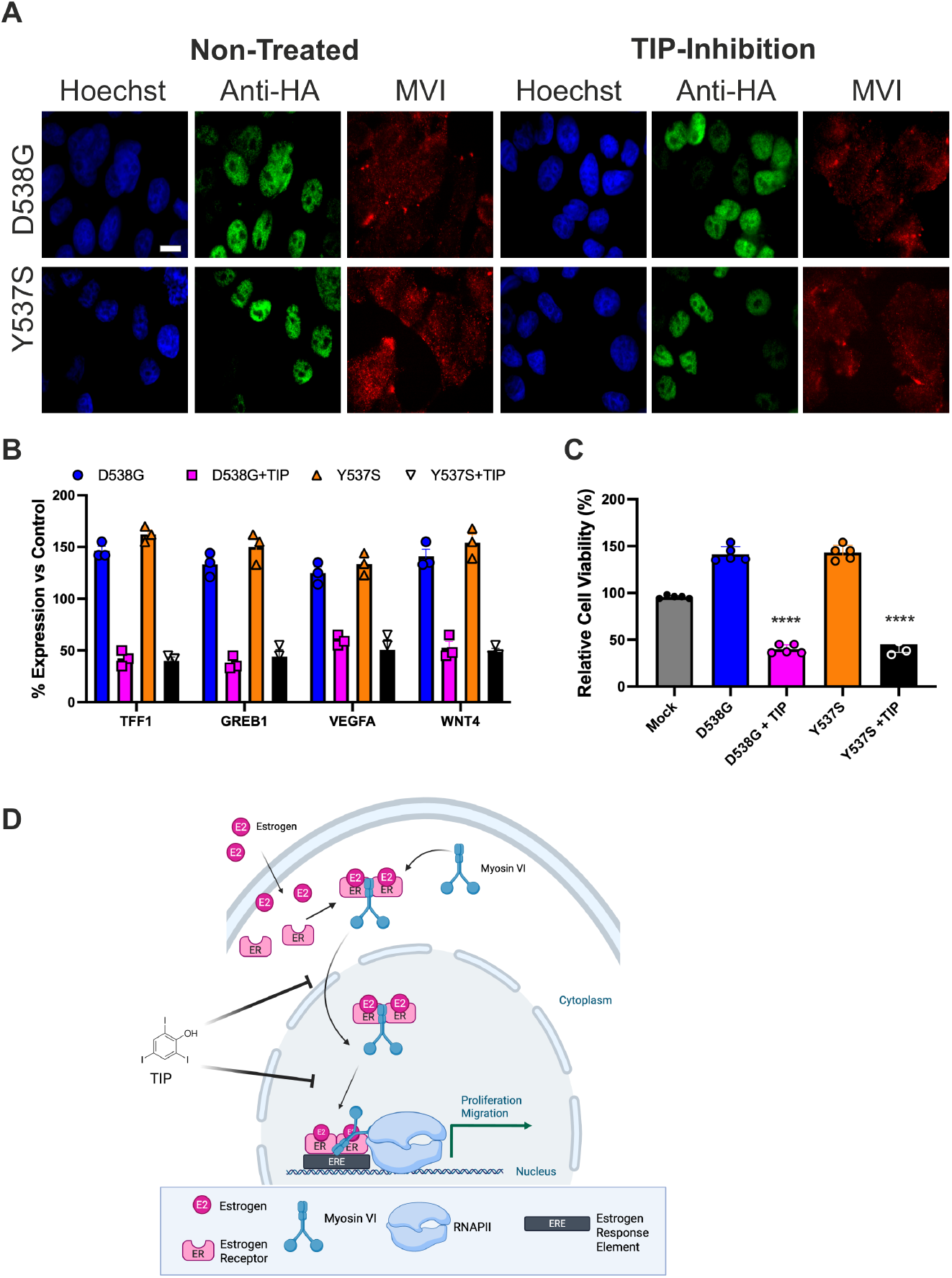
Constitutively active ER mutants are regulated by Myosin VI. (A) Representative widefield Immunofluorescence staining against transfected HA-ER mutant (green), MVI (red) and DNA (blue) in MCF7 cells, in the absence and presence of 25 µM TIP for 6 hrs (Scale bar 10 μm). (B) RT-qPCR gene expression analysis of the stated genes following transfection of ER mutants in the presence and absence of 25 µM TIP for 6 hrs. The plot depicts the percentage change in expression between non-transfected and transfected/treated cells. Error bars represent SEM from three independent experiments. DMSO controls for RT-qPCR are shown in Supplementary Figure 6. All conditions yielded statistically significant changes (p<0.0001) to control by unpaired t-test. (C) MTT cell viability assay under the same conditions as RT-qPCR. All data are normalised to non-treated/non-transfected control. ****p <0.0001 by unpaired t-test compared to D538G or Y537S non-treated condition. (D) Schematic of myosin VI regulation of ER transcription. Based on the data presented here we can propose this model for the interdependency of MVI and the ER clustering. Estrogen binds to the ER which then associates with MVI. The MVI-ER complex is trafficked to the nucleus. The complex then assembles around ERE promoter elements where MVI stabilises interactions with RNAPII to facilitate gene expression. Inhibition of MVI by TIP impacts both nuclear trafficking but also sub-nuclear organisation of MVI-ER at transcription sites, therefore impacting the expression of ER-target genes and the downstream cellular processes, such as cell proliferation and migration.

## DISCUSSION

Our multidisciplinary approach provides new insights into how ER-driven transcription is regulated by MVI, particularly in terms of the ER’s transport and spatial organization within the nucleus. Given that the ER is the key driver of most breast cancers and a major therapeutic target, understanding its partnership with MVI is both novel and clinically relevant.

For two decades, MVI has been linked to transcription ^15,23–26^ and here we have gained further understanding of its nuclear function. MVI is known to stabilise RNAPII by anchoring transcription complexes at genes, and we can now reveal how the ER enables targeting of MVI to specific genes. As previously established, this mechanism enables higher expression levels upon transcription stimulation ^16^.

We propose a model of tight interdependency between MVI and ER (Figure 8D): MVI facilitates the ER’s translocation to the nucleus and organizes ER within sub-nuclear transcription sites, while ER in turn recruits MVI to the nucleus and guides it to specific target genes. This reciprocal relationship could explain why MVI and ER levels correlate so strongly in breast tumour samples. In this manner, the ER recruits MVI as a co-factor to estrogen-responsive promoters, where MVI’s anchoring of RNAPII and local actin networks facilitates the formation of transcription factories ^16^. Two critical findings have arisen from this work: (i) MVI perturbation leads to enhanced ER suppression over monotherapy; (ii) constitutively active ER mutants are still dependent upon MVI.

Our findings suggest that pharmacologically targeting MVI could be a viable strategy to augment current endocrine therapies. In our cell models, combining an MVI inhibitor (TIP) with either a SERM or SERD led to significantly greater suppression of ER target gene expression than either agent alone. This indicates a synergistic effect wherein disrupting MVI’s support of ER function heightens the impact of direct ER antagonism. Importantly, even in the context of hormone-independent ER mutants such as Y537S and D538G, which drive resistance in patients, MVI inhibition reduced transcriptional activity. This highlights MVI as a potential target for ER mutants that no longer respond to hormonal blockade.

While TIP was used here to probe MVI function, it serves as a proof-of-concept. Future efforts could focus on developing MVI inhibitors with suitable potency and specificity for clinical use, potentially opening a new class of co-targeting agents in ER-positive breast cancer therapy. Interestingly, TIP (2,4,6-triiodophenol), also known as AM-24 or Bobel-24, has been used in animal pharmokinetics studies ^27^ and it was shown to induce caspase-independent death in pancreatic carcinoma cell lines ^28^. Therefore, TIP may be more widely applicable in a therapeutic setting. Importantly, our work was conducted in cell culture models; therefore, the efficacy of MVI inhibition in an *in vivo* setting remains to be determined. Future studies should evaluate whether combining MVI inhibitors with endocrine therapy can suppress tumour growth or metastasis in animal models of ER-positive breast cancer.

Our findings with the ER raise the intriguing possibility that MVI might similarly interact with other nuclear receptors (such as the androgen receptor in prostate cancer or progesterone receptor in gynaecological cancers). In fact, there is evidence linking MVI to androgen receptor signalling, though its role in gene expression there is not yet explored ^29^. Moreover, the expression level of MVI within these cancers is also high. It will be worthwhile to investigate whether MVI’s cofactor function is a broader paradigm in hormone-driven cancers.

Consistent with prior reports that MVI supports proliferation and motility in cancer cells ^12,13^, we observed that removing or inhibiting MVI dramatically impairs breast cancer cell growth and migration. Our work extends those observations by pinpointing the ER-driven gene programs that underlie these phenotypes. In other words, MVI’s pro-tumorigenic effects are at least partly due to its facilitation of ER-mediated transcription of genes controlling proliferation and migration. In this manner, the nuclear role of MVI is critical to these cellular functions rather than a direct role connected to the cytoskeleton. This expands upon the intriguing elements of how the roles of cytoplasmic and nuclear myosin proteins can be accurately dissected ^17,30.^ In summary, this study identifies MVI as a central molecular determinant of estrogen receptor dynamics, organisation, and transcriptional output in breast cancer. By coupling mechanical anchoring to transcriptional regulation, MVI integrates nuclear architecture with hormonal signalling to sustain proliferative gene expression. Disruption of MVI not only disassembles ER clustering but also synergises with endocrine therapies and suppresses hormone-independent ER mutants, establishing a new therapeutic axis beyond classical ligand antagonism. These findings redefine MVI as both a mechanistic co-factor and a therapeutic vulnerability in ER-positive tumours. Future efforts to develop selective MVI inhibitors and evaluate their efficacy in pre-clinical models will determine whether targeting this motor can reshape treatment paradigms across hormone-driven cancers.

## ACKNOWLEDGEMENTS

We thank the UKRI-BBSRC (BB/X008460/1), UKRI-MRC (MR/M020606/1), UKRI-STFC (19130001) for funding to C.P.T. Aberration-corrected multi-focal microscopy was performed in collaboration with the Advanced Imaging Center at Janelia Research Campus, a facility jointly supported by the Howard Hughes Medical Institute and the Gordon and Betty Moore Foundation. We also thank Darren Griffin (University of Kent) for sharing of equipment, and Satya Khuon (Janelia Research Campus) for assisting with cell culture. The JF549 dyes were kindly provided by Luke Lavis (Janelia Research Campus).

## AUTHOR CONTRIBUTIONS

C.P.T. conceived the study. Y.H-G., D.L., I. W. S-F., N.F., A.dS., A.S., H.R.K. and C.P.T. designed experiments. N.F., Y.H-G., H.R.K. and C.P.T. designed and cloned constructs. Y.H-G., D.L., I. W. S-F., N.F., A.dS. and A.S. performed imaging experiments on cells and tissues. Y.H-G., N.F., A.dS. and C.P.T. performed single molecule imaging experiments. A.W.C. performed FISH experiments with guidance from P.I.J.E. L.W. and J.A. supported imaging. Y.H-G., N.F. and A.dS. performed and analyzed the genomics experiments. Y.H-G., D.L., I. W.S-F. and A.dS. performed cell phenotypic assays. L.W., J.A., T-L.C. and C.P.T. contributed to single molecule data analysis. C.P.T. supervised the study. C.P.T. wrote the manuscript with comments from all authors.

## MATERIALS AND METHODS

### Constructs

A list of constructs are provided in Supplementary Table 3. Constructs generated in this work are described below. Halo or SNAP tags were used to provide a specific protein labeling strategy for live cells ^31^. The SNAP-ER construct was generated by sub-cloning the ER from into SNAP tag from the pSNAP_f_-C1 plasmid (Addgene 58186). Halo-MVI was generated previously ^25^. Plasmids are from Addgene HA-D538G-ER (49500) and HA-Y537S (49499).

### Cell culture and Transfection

MCF7 (ECACC 86012803) and T47D (ECACC 85102201) cells were maintained in Gibco MEM Alpha medium with GlutaMAX (no nucleosides). ZR-75-1 (ECACC 87012601) cells were maintained in Gibco® Roswell Park Memorial Institute (RPMI) 1640 medium. BT-464 (ATCC HTB-20) cells were maintained in Dulbecco’s Modified Eagle Medium (DMEM) with Glucose and L-Glutamine (Lonza Group, BioWhittaker®, Basel, Switzerland). All media was supplemented with 10% Fetal Bovine Serum (Gibco), 100 units/ml penicillin and 100 µg/ml streptomycin (Gibco). All were growth at 37°C and 5% CO_2_, and passaged at 80% for a maximum of 25 passages.

For the transient expression of HA-ER mutants, MCF7 cells grown on glass coverslips were transfected using Lipofectamine 2000 (Invitrogen), following the manufacturer’s instructions. 48 h after transfection, cells were subjected to nuclear staining using Hoechst 33342 (Thermo Scientific), fixed and analysed or subjected to indirect immunofluorescence (see below).

### Cell Treatments

For MVI knock-down experiments, MCF7 cell monolayers, seeded to 30 – 50 % confluency, were transfected with human myosin VI siRNA duplex (5′GGUUUAGGUGUUAAUGAAGtt-3′) (Ambion 4390824) or AllStars Negative Control siRNA duplex (Qiagen 1027280) at a concentration of 50 nM, using Lipofectamine 2000 (Invitrogen), according to the manufacturer’s guidelines. Cells were fixed or harvested after 48 h for further analysis. The same protocol was followed for knock-down of ER with transfection of siRNA duplex (Ambion 4392420 s4823).

For hormone starvation, MCF7 cell monolayers were cultured in no phenol red MEM media (Gibco) with 5% double charcoal stripped FCS (First Link UK) and 1% Penicillin-Streptomycin for 48 hrs. Hormone stimulation was performed by addition of 10nM of Estradiol for 24 hrs.

TIP (2,4,6-triiodophenol) (Sigma), Tamoxifen citrate (Sigma) and Fulvestrant (Selleckchem) were initially suspended in dimethyl sulfoxide (DMSO, Sigma) to stock concentrations and diluted to indicated concentrations in appropriate medium when applied to cells. Upon drug application, cells were cultured at 37 °C, 5% CO2 for at least 1hr (25 µM TIP) or 6 hr (Tamoxifen/Fulvestrant). Drug replenished at 6-, 24-, and 30-hours where applicable.

### Stable cell line generation

The stable cell lines used in this study are named as MCF7-Halo MVI (pHalo-MVI vector stably expressed in MCF7) and MCF7-SNAP-ER (pSNAP-ER vector stably expressed in MCF7). The MCF7-Halo MVI were generated as described in ^25^. To generate Hela cells stably expressing pSNAP-ER, the plasmid was transfected in 6-well plates using lipofectamine 2000 protocol (Thermo Fisher Scientific). The transfected cells were selected using optimal concentrations of G418 antibiotic (G418 Sulfate, Gibco) in the complete media (0.5mg/ml) for 9-10 days until most of the untransfected cells were dead and those that survived would have integrated the desired plasmid. The cells were harvested when they reached about 60-70% confluency and were expanded into multiple T75 flasks with 1:10 ratio. Some cells at this stage were seeded onto coverslips and stable transfection of desired plasmids was confirmed by using specific fluorescent ligands to SNAP-tag (SNAP-Cell Oregon Green, NEB). The cells seeded in T75 flasks were allowed to grow for further 3-4 weeks in complete media with G418 replaced twice a week. When the cells reached high confluency, they were frozen down as polyclonal stable cell line.

### CRISPR/Cas9 mediated knockout of Myo6 in MCF7 cells

To generate Myo6 knockout MCF7 cells, a sgRNA targeting exon2 (target sequence: 5’ GTTCAATTGTTAAGCTGTCG 3’) was designed using an online CRISPR gRNA design software CHOPCHOP (https://chopchop.cbu.uib.no), and an Alt-R™ CRISPR-Cas9 sgRNA was obtained from Integrated DNA Technologies. The ribonucleoprotein (RNP) complex was pre-assembled by incubating sgRNA and Cas9 protein at a 9:1 molar ratio (180pmol:20pmol) in Lonza nucleofector solution (V4XC-1032) in a total volume of 30ul, followed by incubation for 10 minutes at room temperature. The RNP complex was then electroporated into MCF7 cells using Lonza 4D nucleofector^®^ X-unit according to the manufacturer’s protocol. The electroporated cells were carefully resuspended in 70 µl of prewarmed media and seeded into two wells of a prewarmed 24 well plate. After 72 hrs, DNA was isolated from half of the electroporated cells to assess genome-editing efficiency. The remaining cells were maintained for an additional week before performing serial dilution. Single-cell clones were established in 96 well plates by serial dilution, expanded and screened by western blot.

### Immunofluorescence

Cells were seeded onto 13-mm sterile circular glass (N1.5) coverslips at a density of 160k cell/ml and cultured as above for 24-hours to allow cell attachment. Transfected and non-transfected MCF7 cells were fixed for 15 min at room temperature in 4% (w/v) paraformaldehyde (PFA) in PBS and residual PFA was quenched for 15 min with 50 mM ammonium chloride in PBS. All subsequent steps were performed at room temperature. Cells were permeabilised and simultaneously blocked for 15 min with 0.1 % (v/v) Triton X-100 and 2 % (w/v) BSA in PBS. Cells were then immuno-stained against the endogenous proteins by 1 h incubation with the indicated primary and subsequently the appropriate fluorophore-conjugated secondary antibody (details below), both diluted in 2 % (w/v) BSA in PBS. The following antibodies were used at the indicated dilutions: rabbit anti-myosin VI (1:200, Atlas-Sigma HPA035483), mouse Anti-ERa (1:200, Santa Cruz sc-8002), mouse anti-HA tag (1:300, Abcam ab18181), donkey anti-mouse Alexa Fluor 488-conjugated (1:500, Abcam Ab181289), Donkey anti-rabbit Alexa Fluor 647-conjugated (1:500, Abcam Ab181347). Coverslips were mounted on microscope slides with Mowiol (10% (w/v) Mowiol 4-88, 25% (w/v) glycerol, 0.2 M Tris-HCl, pH 8.5), supplemented with 2.5% (w/v) of the anti-fading reagent DABCO (Sigma).

### Tissue Arrays

Paraffin-embedded human breast tumour tissue arrays (GTX21447, GeneTex) were first baked at 60 °C for 30 minutes as per manufacturer’s instructions. Paraffin dehydration followed through incubations with xylene (10 minutes), xylene (5 minutes), 100% ethanol (10 minutes), 100% ethanol (5 minutes), 95% ethanol (3 minutes), 70% ethanol (3 minutes), 50% ethanol (3 minutes), and finally washed in dH2O. For antigen retrieval, tissues were heated in citrate buffer (10 mM, pH 6.0) for 10 minutes in a microwave at full power, allowed to cool for 20 minutes at room temperature then cooled further under dH2O and once in PBS for 5 minutes. Array area was surrounded with PAP pen (ab2601, Abcam, Cambridge, UK) and tissues simultaneously blocked and permeabilized in 5% BSA/5% FBS/0.3% triton-X/PBS at room temperature for 2 hours. Primary antibody (see above) in 5% BSA/5% FBS/0.01% triton-X/PBS was added to tissue samples and incubated at 4 °C overnight. Following this, samples were left at room temperature for 1 hour, then washed three times in 0.05% Tween/PBS, for 15 minutes each. Secondary antibody (see above) and Hoechst stain were then added to samples in 5%BSA/5%FBS/0.01% triton-X/PBS for 2 hours at room temperature. Samples were then washed three times in 0.05% Tween/PBS and once in dH2O, 15 minutes each. Finally, a coverslip was mounted to the slide with Mowiol/2.5% DABCO and slides left overnight to set at 4 °C. H-Scores were calculated from three independent tissue arrays, as described in ^32^.

### Immunoblot Analysis

The total protein concentration was determined by Bradford Assay (Sigma) following the manufacturer’s instructions. Cell lysates were heat-denatured and resolved by SDS-PAGE. The membrane was probed against the endogenous proteins by incubation with primary Rabbit anti-myosin VI (1:500, Atlas-Sigma HPA035483-100UL), mouse anti-ER (1:200, Santa Cruz sc-8002), rabbit anti-beta-actin (1:5000, Abcam ab8227), rabbit Anti-H3 (1:1000, Cell Signalling Technologies, 9715) and subsequently secondary Goat anti-rabbit antibody (1:15000 Abcam ab6721) or Goat anti-mouse antibody (1:15000, Abcam ab97023) coupled to horseradish peroxidase. The bands were visualised using the ECL Western Blotting Detection Reagents (Invitrogen) and the images were taken using Syngene GBox system. Images were processed in ImageJ.

### Nuclei Isolation

The nuclei isolation protocol was based on modified protocol ^33,34^. MCF7 cells were washed once with ice-cold PBS, then washed in ice-cold Hypotonic Buffer N (10 mM Hepes pH 7.5, 2 mM MgCl_2_, 25 mM KCl supplemented with 1 mM PMSF,1 mM DTT and 1x Halt Protease Inhibitor Cocktail (Thermo Fisher Scientific)). Cells well then re-suspended in ice-cold hypotonic buffer N and incubated for 1 h on ice. Cells were then homogenized on ice with a glass Dounce homogeniser (Wheaton). Cell lysate was supplemented with 2 M sucrose solution and mixed well by inversion before centrifugation. The supernatant which corresponded to the cytoplasmic fraction, was aliquoted and stored at −80 °C. The pellet, which corresponded to isolated nuclei, was further cleaned by washing in ice-cold Buffer N (10 mM Hepes pH 7.5, 2 mM MgCl_2_, 25 mM KCl, 250 mM sucrose, supplemented with 1 mM PMSF, 1 mM DTT and 1x Halt Protease Inhibitor Cocktail). The nuclei pellet was re-suspended either in ice-cold Buffer N and used immediately or in freezing medium (70% glycerol in buffer N), to yield a concentration of 4*10^6^ nuclei ml^−1^. Nuclei were aliquoted on ice and stored at −80 °C.

### Fluorescence Imaging

Cells were visualised using the ZEISS LSM 980 confocal microscope equipped with a Plan-Apochromat 63×/1.4 NA oil immersion objective lens (Carl Zeiss, Cat. No. 420782-9900-000). Three excitation laser lines, 405 nm, 488 nm and 642 nm, were employed to excite the respective fluorophores. Built-in main beam splitters (Carl Zeiss, MBS-405, MBS-488, and MBS-642) directed the laser beams onto the samples.

Emission signals were collected using spectral detection windows of 410–524 nm (Hoechst), 493– 578 nm (green channel), and 564–697 nm (red channel). Detection was achieved using two multi-anode photomultiplier tubes (MA-PMTs) and one gallium arsenide phosphide (GaAsP) detector. The green fluorescence channel was acquired with the GaAsP detector, while blue (Hoechst) and red signals were collected using MA-PMTs. Image acquisition and rendering were performed using ZEN software (Carl Zeiss, ZEN 2.3). All images were then analysed by ImageJ.

### STORM Imaging

Cells were seeded on pre-cleaned No 1.5, 25-mm round glass coverslips, placed in 6-well cell culture dishes. Glass coverslips were cleaned by incubating them for 3 hours, in etch solution, made of 5:1:1 ratio of H_2_O : H_2_O_2_ (50 wt. % in H_2_O, stabilized, Fisher Scientific) : NH_4_OH (ACS reagent, 28-30% NH_3_ basis, Sigma), placed in a 70°C water bath. Cleaned coverslips were repeatedly washed in filtered water and then ethanol, dried and used for cell seeding. Transfected or non-transfected cells were fixed in pre-warmed 4% (w/v) PFA in PBS and residual PFA was quenched for 15 min with 50 mM ammonium chloride in PBS. Immunofluorescence was performed in filtered sterilised PBS. Cells were permeabilized and simultaneously blocked for 30 min with 3% (w/v) BSA in PBS or TBS, supplemented with 0.1 % (v/v) Triton X-100. Permeabilized cells were incubated for 1h with the primary antibody and subsequently the appropriate fluorophore-conjugated secondary antibody, at the desired dilution in 3% (w/v) BSA, 0.1% (v/v) Triton X-100 in PBS or TBS. The antibody dilutions used were the same as for the normal immunofluorescence protocol (see above), except from the secondary antibodies which were used at 1:250 dilution. Following incubation with both primary and secondary antibodies, cells were washed 3 times, for 10 min per wash, with 0.2% (w/v) BSA, 0.05% (v/v) Triton X-100 in PBS or TBS. Cells were further washed in PBS and fixed for a second time with pre-warmed 4% (w/v) PFA in PBS for 10 min. Cells were washed in PBS and stored at 4 °C, in the dark, in 0.02% NaN_3_ in PBS, before proceeding to STORM imaging.

Before imaging, coverslips were assembled into the Attofluor® cell chambers (Invitrogen). Imaging was performed in freshly made STORM buffer consisting of 10 % (w/v) glucose, 10 mM NaCl, 50 mM Tris - pH 8.0, supplemented with 0.1 % (v/v) 2-mercaptoethanol and 0.1 % (v/v) pre-made GLOX solution which was stored at 4 °C for up to a week (5.6 % (w/v) glucose oxidase and 3.4 mg/ml catalase in 50 mM NaCl, 10 mM Tris - pH 8.0). All chemicals were purchased from Sigma.

Imaging was undertaken using the Zeiss Elyra PS.1 system. Illumination was from a HR Diode 642 nm (150 mW) and HR Diode 488 nm (100 mW) lasers where power density on the sample was 7-14 kW/cm^2^ and 7-12 kW/cm^2^, respectively.

Imaging was performed under highly inclined and laminated optical (HILO) illumination to reduce the background fluorescence with a 100x NA 1.46 oil immersion objective lens (Zeiss alpha Plan-Apochromat) with a BP 420-480/BP495-550/LP 650 filter. The final image was projected on an EMCCD camera (Andor iXon^EM+^ DU-897 back-illuminated EMCCD) with 25 msec exposure time per frame for 20000 frames.

Image processing was performed using the Zeiss Zen Black software. Where required, two channel images were aligned following a calibration using a calibration using pre-mounted MultiSpec bead sample (Carl Zeiss, 2076-515). The channel alignment was then performed in the Zeiss Zen Black software using the Affine method to account for lateral, tilting and stretching between the channels. The calibration was performed during each day of measurements.

The images were then processed through our STORM analysis pipeline using the Zen Black software. Single molecule detection and localisation was performed using a 9-pixel mask with a signal to noise ratio of 6 in the “Peak finder” settings while applying the “Account for overlap” function. This function allows multi-object fitting to localise molecules within a dense environment. Molecules were then localised by fitting to a 2D Gaussian.

The render was then subjected to model-based cross-correlation drift correction and detection grouping to remove detections within multiple frames. Typical localisation precision was 20 nm for Alexa-Fluor 647 and 30 nm for Alexa-Fluor 488. The final render was then generated at 10 nm/pixel and displayed in Gauss mode where each localisation is presented as a 2D gaussian with a standard deviation based on its precision. The localisation table was exported as a comma-separated values (csv) file for import into Clus-DoC.

### Clus-DoC

The single molecule positions were exported from Zeiss Zen Black version and imported into the Clus-DoC analysis software ^20^ (https://github.com/PRNicovich/ClusDoC). The region of interest was determined by the nuclear staining. First the Ripley K function was completed on each channel identifying the r max. The r max was then assigned for DBSCAN if one channel was being analysed or Clus-Doc if two channel colcalisation was being analysed. The MinPts was 3 and a cluster required 10 locations, with smoothing set at 7 nm and epsilon set at the mean localization precision for the dye. All other analyses parameters and colocalisation thresholds remained at default settings ^20^. Data concerning each cluster was exported and graphed using Plots of Data.

### Multi-focal Imaging and Particle Tracking Analysis

Cells stably or transiently expressing Halo-tag or SNAP-tag constructs were labelled for 15 min with HaloTag-JF549 or SNAP-tag-JF549 ligand, respectively, in cell culture medium at 37°C, 5% CO_2_. 10 nM ligand was used to label Halo-tagged myosin VI constructs, whereas 50 nM ligand was used to label SNAP-tagged ER. Cells were washed for 3 times with warm cell culture medium and then incubated for further 30 min at 37°C, 5% CO_2_. Cells were then washed three times in pre-warmed FluoroBrite DMEM imaging medium (ThermoFisher Scientific), before proceeding to imaging.

Single molecule imaging was performed using an aberration-corrected multifocal microscope (acMFM), as described by Abrahamsson et al. ^22^. Briefly, samples were imaged using 561nm laser excitation, with typical irradiance of 4-6 kW/cm^2^ at the back aperture of a Nikon 100x 1.4 NA objective. Images were relayed through a custom optical system appended to the detection path of a Nikon Ti microscope with focus stabilization. The acMFM detection path includes a diffractive multifocal grating in a conjugate pupil plane, a chromatic correction grating to reverse the effects of spectral dispersion, and a nine-faceted prism, followed by a final imaging lens.

The acMFM produces nine simultaneous, separated images, each representing successive focal planes in the sample, with ca. 20 µm field of view and nominal axial separation of ca. 400nm between them. The nine-image array is digitized via an electron multiplying charge coupled device (EMCCD) camera (iXon Du897, Andor) at up to 32ms temporal resolution, with typical durations of 30 seconds.

3D+t images of single molecules were reconstructed via a calibration procedure, implemented in Matlab (MathWorks), that calculates and accounts for (1) the inter-plane spacing, (2) affine transformation to correctly align each focal plane in the xy plane with respect to each other, and (3) slight variations in detection efficiency in each plane, typically less than ±5-15% from the mean.

Reconstructed data were then subject to pre-processing, including background subtraction, mild deconvolution (3-5 Richardson-Lucy iterations), and/or Gaussian de-noising prior to 3D particle tracking using the MOSAIC software suite ^35^. Parameters were set where maximum particle displacement was 400 nm and a minimum of 10 frames was required. Tracks were reconstructed, and diffusion constants were extracted via MSD analysis ^36^ using custom Matlab software assuming an anomalous diffusion model.

### 2D Fish Analysis

MCF7 cells were cultured and cells were starved and then stimulated with estradiol, as described above. They were additionally treated with 25 µM TIP for 4 hrs. Once all treatments had been completed cells were washed with PBS and then harvested. The cells were then centrifuged at 500 x g for 10 mins at RT. The cells were resuspended in 75 mM KCl and incubated at 37°C for 20 mins before repeating the centrifugation. The cells were then resuspended in Fix solution (3 parts methanol: 1 part glacial acetic acid) and centrifuged at 500 x g for 10 mins. This process was repeated three times to wash the cells into the Fix solution. A Vysis HYBrite Hybridization System was preheated to 37°C. 10µL of cells in Fix solution were dropped directly onto a microscope slide followed by 10µl of Fix solution only. The slide was left to dry at RT and then immersed into 2x saline-sodium citrate solution (SSC) for 2 mins at RT without agitation. The slides were then dehydrated in an ethanol series (2 mins in 70%, 85% and 100% ethanol) at RT. They were then air dried and placed on the hybridisation system at 37°C. Cover slips were prepared with 10µL of the chromosome paint (Chromosome 2 and 21 Cytocell) and placed onto the slides. The coverslips were sealed using rubber cement and placed back onto the hybridisation system at 75°C for 2 mins. The samples were left within a hybridisation chamber overnight in the dark at 37°C. The rubber cement and coverslips were removed and the slides were immersed into 0.4xSSC pH 7.0 at 72°C for 2 mins, without agitation. The slides were drained and then placed into 2xSSC supplemented with 0.05% Tween-20 at RT for 30 secs. The slides were air-dried and 10µl of DAPI was placed onto each sample and a coverslip was placed and sealed onto the samples. Images were taken using an Olympus BX53 with a 40x oil-immersion objective. Over 60 nuclei per condition were then analysed using the nuclear morphology analysis software, (https://bitbucket.org/bmskinner/nuclear_morphology/wiki/Home) (65).

### Incucyte live cell imaging

Cells were seeded onto 96-well tissue culture dishes at equal densities in 3 replicates. After attachment over-night, cells were transfected with MVI siRNA, or scrambled siRNA (Qiagen). Photomicrographs were taken every hour using an IncuCyte live cell imager (Essen Biosciences, Ann Harbor, MI) and confluency of cultures was measured using IncuCyte software. Confluency values between wells were normalised to initial confluency for comparison.

### Scratch wound-healing assay

Cells were seeded onto 96-well tissue culture dishes at equal densities reaching 100% confluence. A scratch was made at the center of each well using the Incucyte Woundmaker Tool. Photomicrographs were taken every hour using an IncuCyte live cell imager (Essen Biosciences, Ann Harbor, MI) and confluency of cultures was measured using IncuCyte software.

### Colony forming assay

Colony formation was evaluated by seeding approximately 5,000 MCF7 cells and MCF7 MVI knockout cells into 24-well plates containing 1 mL MEM supplemented with 10% FBS and 1% pen/strep. Plates were incubated at 37 °C with 5 % CO_2_ for 1–9 days. Single cells were identified on day 1, and colony growth was monitored every 24 hours using an EVOS M7000 imaging system (TransLight channel, 4× objective, 12 bits per pixel, 1×1 binning). Colonies were counted as particles in ImageJ/Fiji.

### Invasion/Migration assay

Measurements were performed using the CHEMICON cell invasion assay 24 well system (Sigma) with 8 µm pores. Well inserts were washed with serum free media before plating MCF7 cells. Media with 10% FBS was added to the lower chamber. Cells were incubated for 24 hours at 37°C. Cells were removed from inside the insert before incubating with cell detachment solution for 30 min (90131). Cells were then incubated with cell lysis buffer (90130) and CyQaunt GR Dye (90132) (1:75 dilution) for 15 min. Equal volumes of the solute were transferred to a 96 well plate. The fluorescence at 480/520 was measured for each sample.

### MTT Assay (Cytotoxicity/Viability Assay)

Cell viability was assessed after drug treatments using the 3-(4,5-dimethylthiazol-2-yl)-2,5-diphenyltetrazolium bromide (MTT) assay. Cells were seeded into 96 well plates and treatments were performed for 24 hrs. 10 µl MTT stock (Biotium BT30006)) was added to the culture medium and cells were incubated at 37°C with 5 % CO2 for 4 hrs. Next, the cells were incubated with 200 μl of dimethyl sulfoxide (DMSO; Sigma) for an additional 5 min, and their viability was measured using a microplate reader at 570 nm. The MTT assays were performed at least three times for each concentration of drug and the percentage of surviving cells relative to the non-treated control was determined.

### RNA extraction and RT-qPCR

RNA from MCF7 cells was extracted using Gene Jet RNA purification kit (Thermo scientific) according to manufacturer’s protocol. The RNA concentration was measured using Geneflow Nanophotometer and RT-qPCR was performed with one-step QuantiFast SYBR Green qPCR kit (Qiagen) using 50ng of RNA in each sample. A list of qPCR primers is given in Supplementary Table 4.

### RNA-seq and analysis

Total RNA was extracted from three replicates of WT, MVI KD and Scrambled siRNA. Ice cold TRIzol reagent was added to each culture and homogenised. The mixture was then incubated for 5 mins at room temperature then chloroform was added to the lysis and incubated for 3 mins. The samples were then centrifuged at 12,000 x g at 4°C. The colourless aqueous phase was collected. The RNA was then precipitated with incubation for 10 mins with isopropanol before centrifugation for a further 10 mins at 12,000 x g at 4 °C. The pellet was washed in 75% (v/v) ethanol, vortexed and centrifuged for 5 mins at 7500 x g at 4°C. The RNA pellet is air dried for 10 mins. The pellet is then resuspended in 50µL of RNase-free water containing 0.1mM EDTA and incubated at 55°C for 15 mins to allow the RNA to dissolve. The RNA was then quantified using then 260nm absorbance, ensuring the A260/A280 ratio was approximately 2, therefore implying the sample is pure. The sample was then further purified using the RNeasy kit (Qiagen) where the manufacturer’s protocol was followed exactly. Once the purity and stability had been measured the RNA was then stored at −80°C.

The following procedures were performed by Glasgow Polyomics. The RNA-seq libraries were prepared using Poly-A selection. Resulting libraries concentration, size distribution and quality were assessed on a Qubit fluorometer and on an Agilent 2100 bioanalyzer. Paired-end sequencing (2 × 75 bp) on a HiSeq sequencer.

Sequence reads were trimmed to remove possible adapter sequences and nucleotides with poor quality using Trimmomatic v.0.36. The trimmed reads were mapped to the Homo sapiens GRCh38 reference genome available on ENSEMBL using the STAR aligner v.2.5.2b. BAM files were generated as a result of this step. Unique gene hit counts were calculated by using featureCounts from the Subread package v.1.5.2. After extraction of gene hit counts, the gene hit counts table was used for downstream differential expression analysis. Using DESeq2, a comparison of gene expression between the customer-defined groups of samples was performed. The Wald test was used to generate *p* values and log2 fold changes. Genes with an adjusted *p* value < 0.05 and absolute log2 fold change >1 were called as differentially expressed genes for each comparison.

Differentially expressed genes by at least 2-fold log2(FC)≥1 and adjusted p-value of <0.05 for upregulated genes and log2(FC)≤-1 and adjusted p-value of <0.05 for downregulated genes) between the samples which were then subjected to Gene Ontology (GO) enrichment using ShinyGo v0.60 (http://bioinformatics.sdstate.edu/go/) ^37^. RNA-Seq data were deposited in the Gene Expression Omnibus (GEO) database under the accession number GSE300165.

### Chromatin Immunoprecipitation (ChIP)

To identify specific MVI-DNA interactions, ChIP was performed using rabbit anti-myosin VI (Atlas-Sigma Sigma HPA035483-100UL). A confluent T175 flask (10×10^6^ – 30×10^6^) of MCF7 cells was crosslinked by adding formaldehyde dropwise directly to the media to a final concentration of 0.75% and was left for gentle rotation at room temperature (RT) for 10min. To stop the reaction, glycine was added to a final concentration of 125 mM and was incubated with shaking for 5 min at RT. The cells were washed twice with 10 ml of cold PBS and were scraped in 5-8 ml of cold PBS. All cells were collected and centrifuged at 1000xg, 4°C for 5 min. The pellet was re-suspended in ChIP lysis buffer (750 μL per 1×10^7^ cells) (lysis buffer: 50 mM HEPES-KOH pH 7.5, 140 mM NaCl, 1 mM EDTA pH8, 1 % TritonX-100, 0.1 % Sodium Deoxycholate, 0.1 % SDS and Protease Inhibitors) and was incubated on ice for 10 min. The cells were sonicated using the diagenode bioruptor sonicator in order to shear DNA to an average fragment size of 200-800 bp. The fragment size was analysed on a 1.5 % agarose gel. After sonication, cell debris was removed by centrifugation for 10 min, 4°C, 8000xg and the supernatant (chromatin) was used for the immunoprecipitation. The sonicated chromatin was snap frozen on dry ice and was stored at −80°C until further use (max storage 3 months).

The chromatin prepared above was diluted 1:10 with RIPA buffer (50 mM Tris-HCl pH8, 150 mM NaCl, 2 mM EDTA pH8, 1% NP-40, 0.5% Sodium Deoxycholate, 0.1% SDS and Protease Inhibitors) and was distributed into 6 tubes (approximately 1×10^6^ cells per IP) - 3 samples for specific antibody (MVI) and 3 samples for the no antibody control (beads only). 10% of diluted chromatin was removed to serve as input sample and was stored at −20°C until further use. All chromatin samples were pre-cleared using the protein A magnetic beads (Thermo Fisher Scientific) for 30 min after which 20 µl of rabbit anti-myosin VI, was added to each of the triplicate Ab samples (1 in 50 dilution) and the tubes were rotated at 4°C, overnight. Next day, 40 µl of protein A magnetic beads (washed three times in RIPA buffer) were added to each of the samples including the no antibody control tubes and were put on rotation at 4°C for 1 h. After 1h, the beads were collected using a magnetic rack and were washed twice in low salt buffer (0.1 % SDS, 1 % Triton X-100, 2 mM EDTA, 20 mM Tris-HCl pH 8, 150 mM NaCl), once in high salt buffer (0.1 % SDS, 1 % Triton X-100, 2 mM EDTA, 20 mM Tris-HCl pH 8, 500 mM NaCl), once in LiCl wash buffer (0.25 M LiCl, 1 % NP-40, 1 % Sodium Deoxycholate, 1 mM EDTA, 10 mM Tris-HCl pH 8) and finally in TE (10mM Tris pH8, 1mM EDTA). DNA was eluted by adding 120 µl of elution buffer (1 % SDS, 100 mM NaHCO_3_) to the beads and vortexing them slowly for 15 min at 30°C. To reverse crosslink the protein-DNA complexes, 4.8 µl of 5M NaCL and 2 µl RNase A (10mg/ml) was added to the elutes including the input sample that was stored at −20°C and they were incubated while shaking at 65°C overnight followed by proteinase K treatment at 60°C for 1 h. The DNA was then purified using phenol:chloroform extraction and the samples were analysed by qPCR using primers in Supplementary Table 4.

### Graphics

Unless stated, data fitting and plotting was performed using Plots of data ^38^, Graphpad Prism and Grafit Version 5 (Erithacus Software Ltd). Cartoons were generated using the BioRender software.

## Data Availability

The data supporting the findings of this study are available from the corresponding author on request.

## REFERENCES

1 Lower, E. E., Glass, E. L., Bradley, D. A., Blau, R. & Heffelfinger, S. Impact of metastatic estrogen receptor and progesterone receptor status on survival. Breast Cancer Res Treat 90, 65–70 (2005). 10.1007/s10549-004-2756-z

2 Yersal, O. & Barutca, S. Biological subtypes of breast cancer: Prognostic and therapeutic implications. World J Clin Oncol 5, 412–424 (2014). 10.5306/wjco.v5.i3.412

3 Manavathi, B., Samanthapudi, V. S. & Gajulapalli, V. N. Estrogen receptor coregulators and pioneer factors: the orchestrators of mammary gland cell fate and development. Front Cell Dev Biol 2, 34 (2014). 10.3389/fcell.2014.00034

4 Lumachi, F., Brunello, A., Maruzzo, M., Basso, U. & Basso, S. M. Treatment of estrogen receptor-positive breast cancer. Curr Med Chem 20, 596–604 (2013). 10.2174/092986713804999303

5 Connor, C. E. et al. Circumventing tamoxifen resistance in breast cancers using antiestrogens that induce unique conformational changes in the estrogen receptor. Cancer Res 61, 2917–2922 (2001).

6 Pearce, S. T., Liu, H. & Jordan, V. C. Modulation of estrogen receptor alpha function and stability by tamoxifen and a critical amino acid (Asp-538) in helix 12. J Biol Chem 278, 7630–7638 (2003). 10.1074/jbc.M211129200

7 McDonnell, D. P., Wardell, S. E. & Norris, J. D. Oral Selective Estrogen Receptor Downregulators (SERDs), a Breakthrough Endocrine Therapy for Breast Cancer. J Med Chem 58, 4883–4887 (2015). 10.1021/acs.jmedchem.5b00760

8 Schneider, R., Barakat, A., Pippen, J. & Osborne, C. Aromatase inhibitors in the treatment of breast cancer in postmenopausal female patients: an update. Breast Cancer (Dove Med Press) 3, 113–125 (2011). 10.2147/BCTT.S22905

9 Huang, D., Yang, F., Wang, Y. & Guan, X. Mechanisms of resistance to selective estrogen receptor down-regulator in metastatic breast cancer. Biochim Biophys Acta Rev Cancer 1868, 148–156 (2017). 10.1016/j.bbcan.2017.03.008

10 Kuukasjarvi, T., Kononen, J., Helin, H., Holli, K. & Isola, J. Loss of estrogen receptor in recurrent breast cancer is associated with poor response to endocrine therapy. J Clin Oncol 14, 2584–2589 (1996). 10.1200/JCO.1996.14.9.2584

11 Magnani, L. et al. Genome-wide reprogramming of the chromatin landscape underlies endocrine therapy resistance in breast cancer. Proc Natl Acad Sci U S A 110, E1490–1499 (2013). 10.1073/pnas.1219992110

12 Dunn, T. A. et al. A novel role of myosin VI in human prostate cancer. Am J Pathol 169, 1843–1854 (2006). 10.2353/ajpath.2006.060316

13 Wang, H., Wang, B., Zhu, W. & Yang, Z. Lentivirus-Mediated Knockdown of Myosin VI Inhibits Cell Proliferation of Breast Cancer Cell. Cancer Biother Radiopharm 30, 330–335 (2015). 10.1089/cbr.2014.1759

14 Fili, N. & Toseland, C. P. Unconventional Myosins: How Regulation Meets Function. Int J Mol Sci 21 (2019). 10.3390/ijms21010067

15 Fili, N. et al. NDP52 activates nuclear myosin VI to enhance RNA polymerase II transcription. Nat Commun 8, 1871 (2017). 10.1038/s41467-017-02050-w

16 Hari-Gupta, Y. et al. Myosin VI regulates the spatial organisation of mammalian transcription initiation. Nat Commun 13, 1346 (2022). 10.1038/s41467-022-28962-w

17 Shahid-Fuente, I. W. & Toseland, C. P. Myosin in chromosome organisation and gene expression. Biochem Soc Trans 51, 1023–1034 (2023). 10.1042/BST20220939

18 Tang, Z. et al. GEPIA: a web server for cancer and normal gene expression profiling and interactive analyses. Nucleic Acids Res 45, W98–W102 (2017). 10.1093/nar/gkx247

19 Dos Santos, A., Gough, R. E., Wang, L. & Toseland, C. P. Measuring Nuclear Organization of Proteins with STORM Imaging and Cluster Analysis. Methods Mol Biol 2476, 293–309 (2022). 10.1007/978-1-0716-2221-6_20

20 Pageon, S. V., Nicovich, P. R., Mollazade, M., Tabarin, T. & Gaus, K. Clus-DoC: a combined cluster detection and colocalization analysis for single-molecule localization microscopy data. Mol Biol Cell 27, 3627–3636 (2016). 10.1091/mbc.E16-07-0478

21 Malkusch, S. et al. Coordinate-based colocalization analysis of single-molecule localization microscopy data. Histochem Cell Biol 137, 1–10 (2012). 10.1007/s00418-011-0880-5

22 Abrahamsson, S. et al. Fast multicolor 3D imaging using aberration-corrected multifocus microscopy. Nat Methods 10, 60–63 (2013). 10.1038/nmeth.2277

23 Cook, A., Hari-Gupta, Y. & Toseland, C. P. Application of the SSB biosensor to study in vitro transcription. Biochemical and biophysical research communications 496, 820–825 (2018). 10.1016/j.bbrc.2018.01.147

24 Fili, N. et al. Competition between two high- and low-affinity protein-binding sites in myosin VI controls its cellular function. The Journal of biological chemistry 295, 337–347 (2020). 10.1074/jbc.RA119.010142

25 Große-Berkenbusch, A. et al. Myosin VI moves on nuclear actin filaments and supports long-range chromatin rearrangements. bioRxiv, 2020.2004.2003.023614 (2020). 10.1101/2020.04.03.023614

26 Vreugde, S. et al. Nuclear myosin VI enhances RNA polymerase II-dependent transcription. Mol Cell 23, 749–755 (2006). 10.1016/j.molcel.2006.07.005

27 Garcia-Capdevila, L. et al. High-performance liquid chromatography analysis of Bobel-24 in biological samples for pharmacokinetic, metabolic and tissue distribution studies. J Chromatogr B Biomed Sci Appl 708, 169–175 (1998). 10.1016/s0378-4347(97)00645-2

28 Parreno, M. et al. Bobel-24 and derivatives induce caspase-independent death in pancreatic cancer regardless of apoptotic resistance. Cancer Res 68, 6313–6323 (2008). 10.1158/0008-5472.CAN-08-1054

29 Loikkanen, I. et al. Myosin VI is a modulator of androgen-dependent gene expression. Oncol Rep 22, 991–995 (2009).

30 Gawor, M., Lehka, L., Lambert, D. & Toseland, C. P. Actin from within - how nuclear myosins and actin regulate nuclear architecture and mechanics. J Cell Sci 138 (2025). 10.1242/jcs.263550

31 Toseland, C. P. Fluorescent labeling and modification of proteins. J Chem Biol 6, 85–95 (2013). 10.1007/s12154-013-0094-5

32 Ma, H. et al. Quantitative measures of estrogen receptor expression in relation to breast cancer-specific mortality risk among white women and black women. Breast Cancer Res 15, R90 (2013). 10.1186/bcr3486

33 Dos Santos, A. et al. DNA damage alters nuclear mechanics through chromatin reorganization. Nucleic Acids Res (2020). 10.1093/nar/gkaa1202

34 Lherbette, M. et al. Atomic Force Microscopy micro-rheology reveals large structural inhomogeneities in single cell-nuclei. Sci Rep 7, 8116 (2017). 10.1038/s41598-017-08517-6

35 Sbalzarini, I. F. & Koumoutsakos, P. Feature point tracking and trajectory analysis for video imaging in cell biology. J Struct Biol 151, 182–195 (2005). 10.1016/j.jsb.2005.06.002

36 Aaron, J., Wait, E., DeSantis, M. & Chew, T. L. Practical Considerations in Particle and Object Tracking and Analysis. Curr Protoc Cell Biol 83, e88 (2019). 10.1002/cpcb.88

37 Ge, S. X., Jung, D. & Yao, R. ShinyGO: a graphical enrichment tool for animals and plants. Bioinformatics (2019). 10.1093/bioinformatics/btz931

38 Postma, M. & Goedhart, J. PlotsOfData-A web app for visualizing data together with their summaries. PLoS biology 17, e3000202 (2019). 10.1371/journal.pbio.3000202

