## Supplementary Data for "Myosin VI orchestrates estrogen-driven gene expression in breast cancer cells"

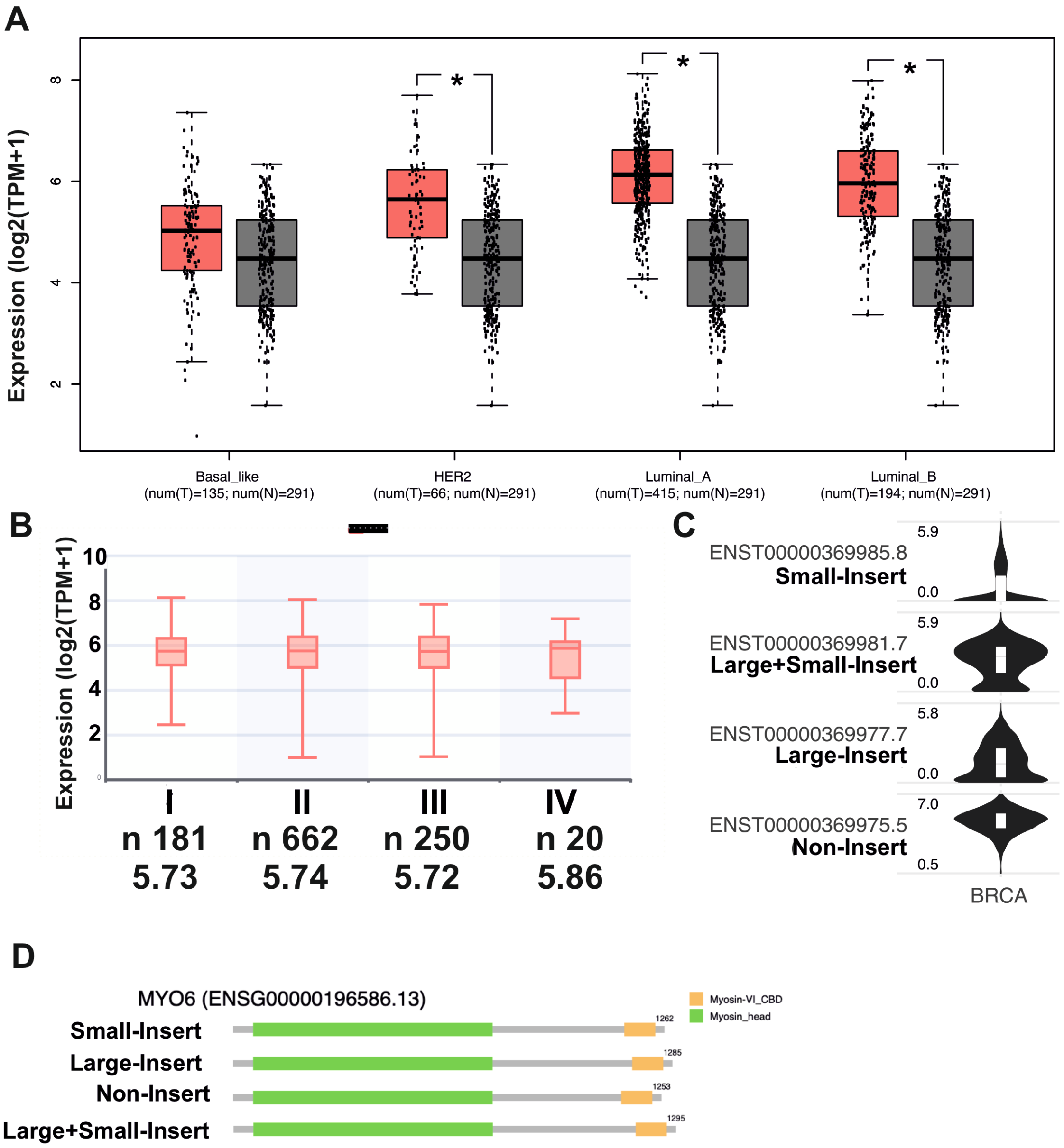


**Supplementary Figure 1: Expression of MYO6 in breast cancer.**
(A) Expression level of *MYO6* in breast cancer vs normal breast tissue separated by breast cancer classification. (B) Expression level (Transcripts per million) of *MYO6* in breast cancer classified by disease stage. (C) The violin-plots show the expression level (log2(TPM+1) of each MYO6 isoform. The isoform structure is shown in (D).

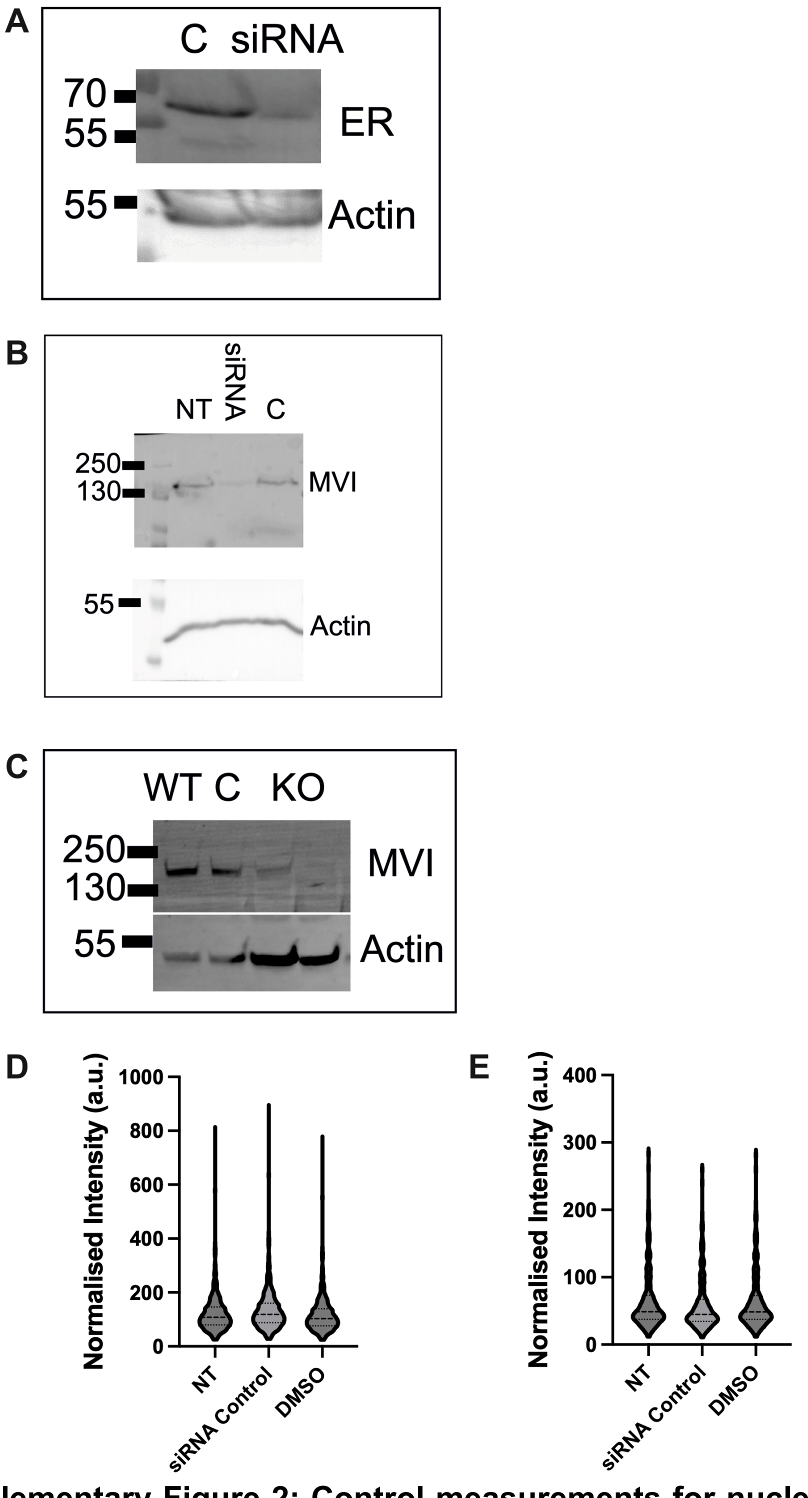


**Supplementary Figure 2: Control measurements for nuclear localisation of ER and MVI.**

(A) Western blots for the ER knockdown in MCF7 cells 48 hrs after transfection. (B) Western blots for the MVI knockdown in MCF7 cells 48 hrs after transfection. (C) Western blots for the MVI knockout MCF7 cells. (D) and (E) Quantification of nuclear MVI and ER normalised intensity across the experimental conditions. n=560 cells per condition across three independent experiments. WT: Wild type; NT: Non-treated; KO: MVI knockout.

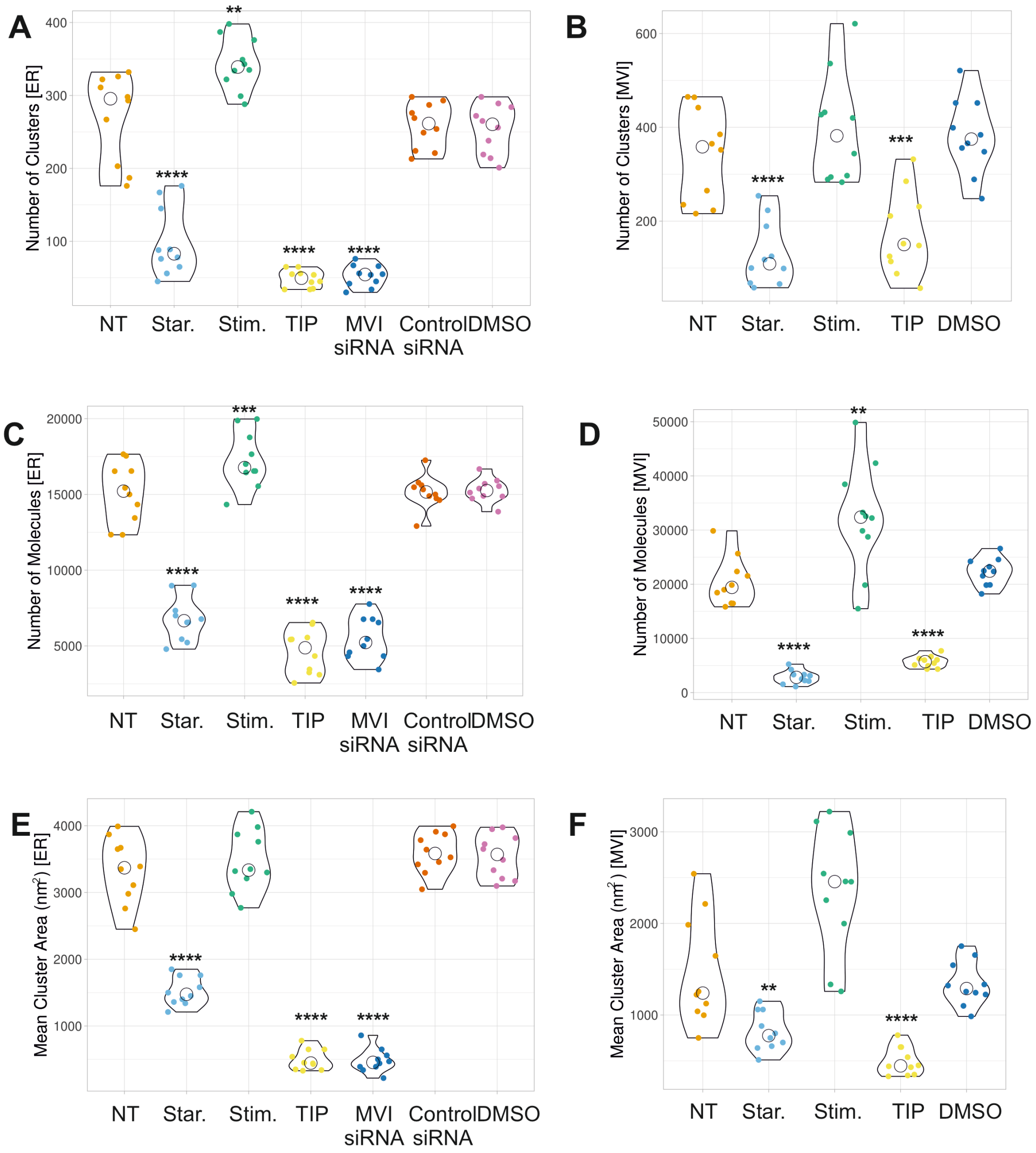


**Supplementary Figure 3: Cluster analysis of nuclear ER and MVI.**
MCF7 cells were stained for MVI and ER under non-treated, hormone starvation, estradiol stimulation, TIP and MVI knockdown (siRNA MVI) conditions (scale bar 2 □m). Hormone starvation occurred for 48 hrs and stimulation was performed upon starved cells for 24 hr with 10 nM estradiol. siRNA knockdowns were performed for 48 hrs. 25 μM TIP for 6 hrs.
Clusters are defined by detecting a minimum of 5 molecules within a search area corresponding to the STORM localisation precision. The search area then propagates and a group of molecules is considered to be a cluster if at least 10 molecules are found. Individual data points correspond to the average value for a cell ROI (n = 10). (A) and (B)The values represent the number of clusters from the ROIs for each condition for ER and MVI, respectively. (C) and (D) represents the number of molecules of ER and MVI within the ROI, respectively. (E) and (F) represents the mean cluster area per ROI for ER and MVI, respectively. Only statistically significant

changes are highlighted **p <0.01, ***p <0.001, ****p <0.0001 by unpaired t-test compared to normal conditions.

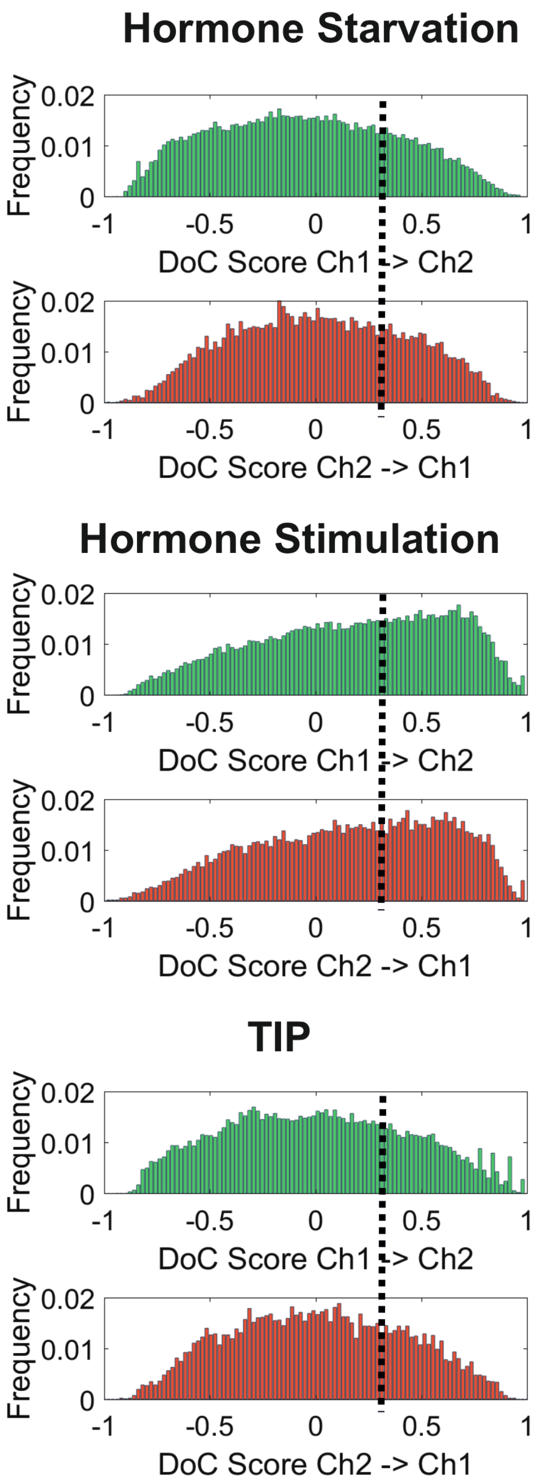


**Supplementary Figure. 4 Colocalisation of MVI and ER.** Representative DoC value histograms for MVI colocalised with ER and ER colocalised with MVI under hormone starvation (48 hrs), hormone stimulation with 10 nM estradiol following starvation, and treatment with 25 μM TIP for 6 hrs. The DoC values are calculated using coordinate-based colocalization analysis (20) in Clus-DoC (19). -1 is segregated, 0 is random distribution and +1 is colocalised. A threshold is applied at values about +0.4 (dashed line) to determine colocalised molecules, as described in (19).

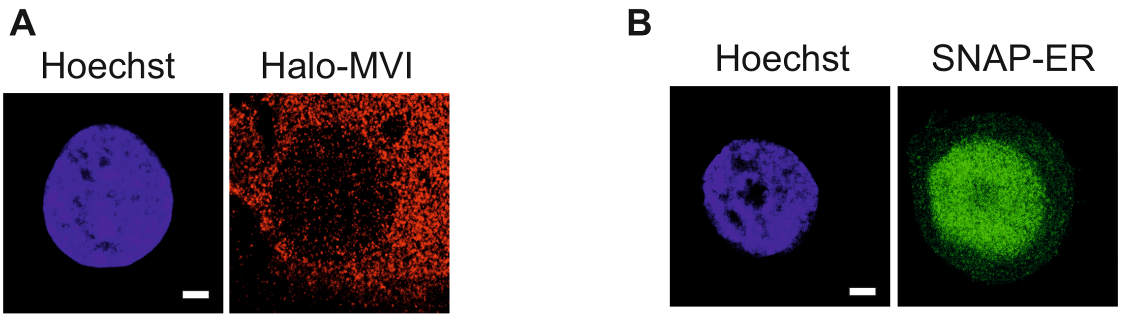


**Supplementary Figure 5: Stable MCF7 cell lines.**
(A) Stable MCF7 Halo-MVI cells stained with Halo-dye TMR. (B) Stable MCF7 SNAP-ER cells stained with SNAP-dye Oregon Green. Scale bar 2 µm.

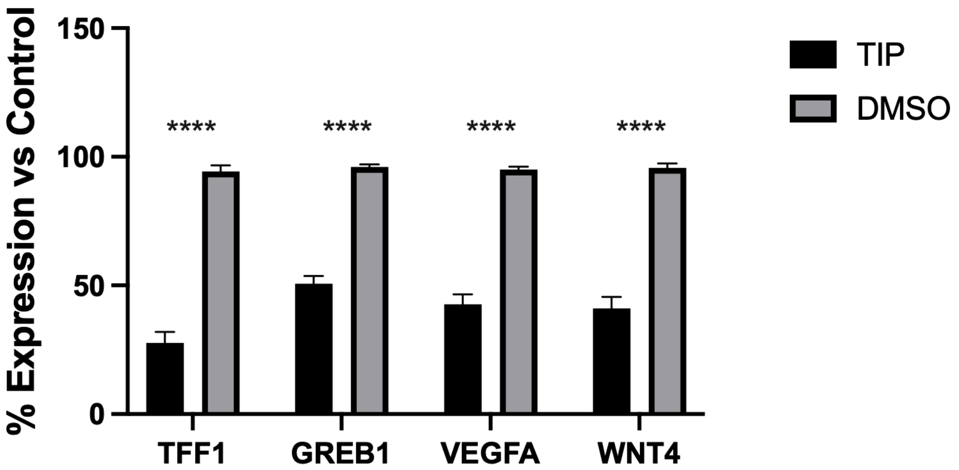


**Supplementary Figure 6: RT-qPCR gene expression.**
RT-qPCR gene expression analysis of the stated genes within MCF7 cells. Cells were treated with 25 μM TIP or DMSO for 6 hrs. The plot depicts the percentage change in expression between non-treated and treated conditions. Error bars represent SEM from three independent experiments. ****p <0.0001 by unpaired t-test compared to normal conditions.

**A**

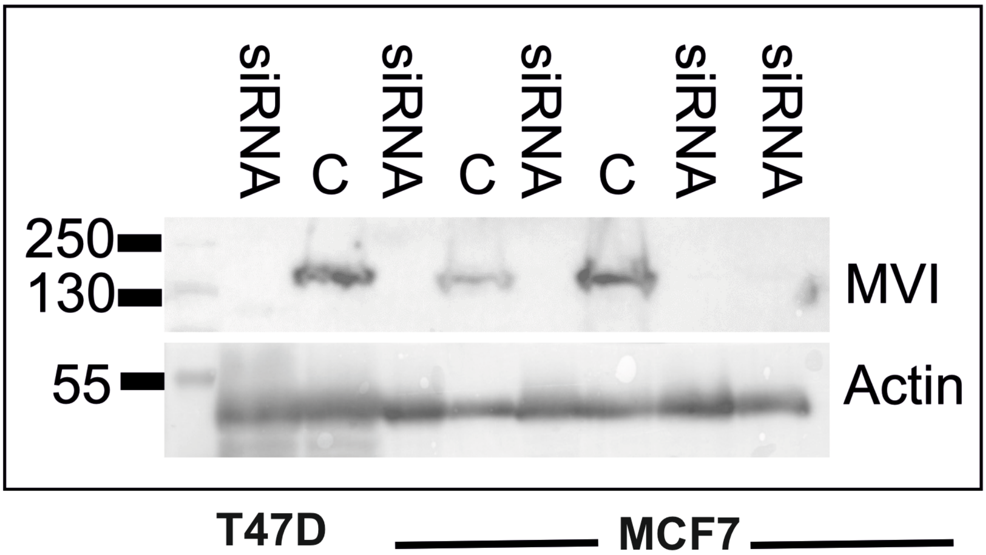


**B**

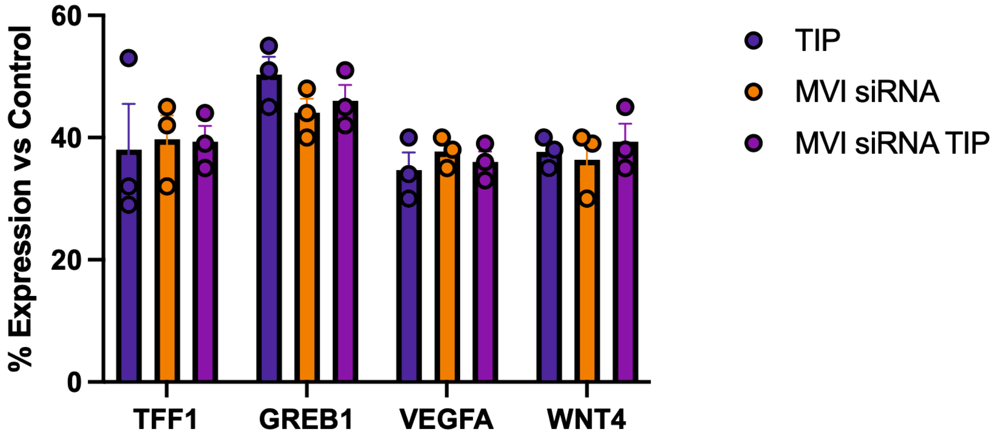


**Supplementary Figure 7: Control western blots for cell-based assays.**
(A) Western blots for the MVI knockdown in T47D and MCF7 cells 48 hrs after transfection. (B) RT-qPCR gene expression analysis of the stated genes. MCF7 cells were treated with 25 μM TIP for 6 hrs, subjected to siRNA knockdown for 24 hrs. or a combined treatment. The plot depicts the percentage change in expression between non-treated and treated cells. Error bars represent SEM from three independent experiments.

A

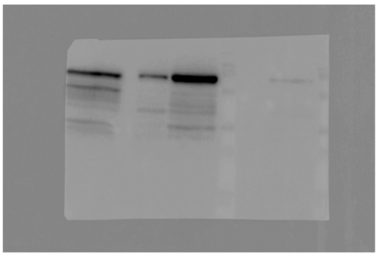

ER Nuclear Isolation +/- E2

B

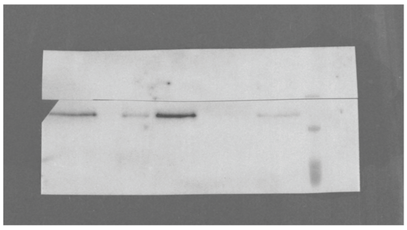

MVI Nuclei Isolation +/- E2

C

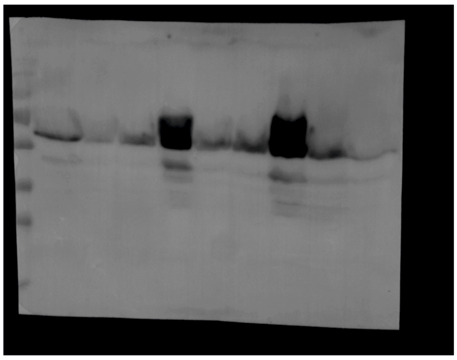

ER siRNA

D

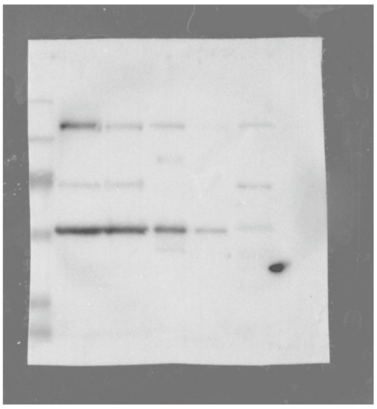

MVI siRNA RNA Seq

E

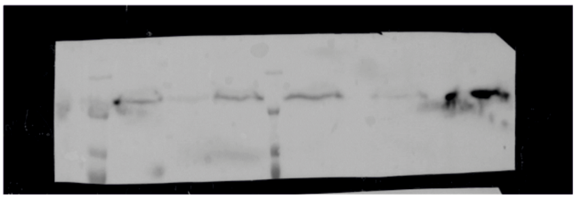

MVI siRNA Imaging

F

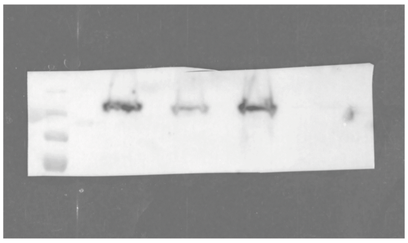

Cell-based assays

**Supplementary Figure 8: Full images for western blots used in this manuscript.** Western blots for the MVI knockdown in T47D and MCF7 cells 48 hrs after transfection.

**Supplementary Table 1. GO Gene Expression analysis: Biological Process**

| nGenes | Pathway Genes | Fold Enrichment | Pathway |
|---|---|---|---|
| 76 | 1086 | 2.270058125 | GO:0009887 Animal organ morphogenesis |
| 80 | 1155 | 2.246783435 | GO:0035295 Tube development |
| 80 | 1192 | 2.177042674 | GO:0072359 Circulatory system development |
| 130 | 2165 | 1.947774439 | GO:0009888 Tissue development |
| 111 | 1859 | 1.93685362 | GO:0022008 Neurogenesis |
| 95 | 1604 | 1.921199442 | GO:0048699 Generation of neurons |
| 97 | 1655 | 1.90119624 | GO:0016477 Cell migration |
| 122 | 2185 | 1.811179942 | GO:0008283 Cell population proliferation |
| 118 | 2120 | 1.80550775 | GO:0006915 Apoptotic proc. |
| 154 | 2868 | 1.741785955 | GO:0009653 Anatomical structure morphogenesis |
| 143 | 2764 | 1.678228953 | GO:0007399 Nervous system development |
| 167 | 3229 | 1.677651064 | GO:0048513 Animal organ development |
| 221 | 4287 | 1.672214561 | GO:0048731 System development |
| 167 | 3304 | 1.639568791 | GO:0009966 Reg. of signal transduction |
| 162 | 3241 | 1.621396361 | GO:0035556 Intracellular signal transduction |
| 185 | 3753 | 1.598992308 | GO:0023051 Reg. of signaling |
| 185 | 3758 | 1.596864857 | GO:0010646 Reg. of cell communication |
| 227 | 4782 | 1.539818368 | GO:0030154 Cell differentiation |
| 227 | 4783 | 1.539496432 | GO:0048869 Cellular developmental proc. |
| 187 | 3988 | 1.521036611 | GO:0009893 Pos. reg. of metabolic proc. |

**Supplementary Table 2. GO Gene Expression analysis: Biological Pathway**

| nGenes | Pathway Genes | Fold Enrichment | Pathway |
|---|---|---|---|
| 17 | 75 | 7.352598791 | Path:hsa04115 p53 signaling pathway |
| 5 | 29 | 5.592747559 | Path:hsa04392 Hippo signaling pathway-multiple species |
| 5 | 32 | 5.068427476 | Path:hsa04215 Apoptosis-multiple species |
| 6 | 41 | 4.747015002 | Path:hsa05219 Bladder cancer |
| 10 | 75 | 4.325058113 | Path:hsa05214 Glioma |
| 11 | 87 | 4.10134821 | Path:hsa05210 Colorectal cancer |
| 9 | 75 | 3.892552301 | Path:hsa01524 Platinum drug resistance |
| 11 | 92 | 3.878448851 | Path:hsa04912 GnRH signaling pathway |
| 16 | 137 | 3.788372069 | Path:hsa04210 Apoptosis |
| 14 | 132 | 3.440387135 | Path:hsa04068 FoxO signaling pathway |
| 14 | 137 | 3.314825561 | Path:hsa04371 Apelin signaling pathway |
| 9 | 96 | 3.041056485 | Path:hsa01522 Endocrine resistance |
| 9 | 96 | 3.041056485 | Path:hsa04713 Circadian entrainment |
| 13 | 153 | 2.756164483 | Path:hsa04921 Oxytocin signaling pathway |
| 16 | 203 | 2.556684599 | Path:hsa05205 Proteoglycans in cancer |
| 14 | 182 | 2.495225834 | Path:hsa04360 Axon guidance |
| 37 | 529 | 2.268815929 | Path:hsa05200 Pathways in cancer |
| 16 | 232 | 2.237099024 | Path:hsa04820 Cytoskeleton in muscle cells |
| 19 | 300 | 2.054402603 | Path:hsa04010 MAPK signaling pathway |
| 80 | 1556 | 1.667760198 | Path:hsa01100 Metabolic pathways |

**Supplementary Table 3. Recombinant DNA.**

| Construct | Source |
|---|---|
| pSNAP-C1 | Addgene 58186 |
| pSNAP-ESR1 | This Study |
| HA-D538G-ER | Addgene 49500 |
| HA-Y537S | Addgene 49499 |

### Supplementary Table 4. Primers for qPCR.

| Sequence | Use |
|---|---|
| GCCACGTCTCCACACATCAG | MYC For |
| TCTTGGCAGCAGGATAGTCCT | MYC Rev |
| CCATGTTGGCCAGGCTAGTC | TFF1 For |
| ACAACAGTGGCTCACGGGGT | TFF1 Rev |
| GCAGTGAAAAAAAGTGTGGCAACTGGG | GREB1 For |
| GACCCACAGAAATGAAAAGGCAGCAAACT | GREB1 Rev |
| GACCTGGCCCAGATAGATCA | NRIP For |
| TATAATGTAGGGGCGCAACC | NRIP Rev |
| GCTGGGTCCTCTGGCTGTTC | RARA For |
| CCGGGATAAAGCCACTCCAA | RARA Rev |
| ATCGCCAGGCCTCCTCACTTGG | VEGFA For |
| TGAGGGAGGCTCCTTCCTC | VEGFA Rev |
| CGCCCGCCTCCGCCGCCAC | WNT4 For |
| CGCAGGGACCGCAGGCACGAA | WNT4 Rev |
| CATGGAGAACAAGGTGATCTG | TFF1 RT-qPCR For. |
| CACTGTACACGTCTCTGTCTG | TFF1 RT-qPCR Rev. |
| ATGGGAAATTCTTACGCTGGAC | GREB1 RT-qPCR For |
| CACTCGGCTACCACCTTCT | GREB1 RT-qPCR Rev |
| TTGCCTTGCTGCTCTACCTCCA | VEGFA RT-qPCR For |
| GATGGCAGTAGCTGCGCTGATA | VEGFA RT-qPCR Rev |
| GCTGGAGAAGTGCGGCTGTGA | WNT4 RT-qPCR For |
| CCACAAACGACTGTGAGAAGGC | WNT4 RT-qPCR Rev |
